# Chemotherapy induces a YAP1-dependent fetal conversion to human Colorectal Cancer cells that is predictive of poor patient outcome

**DOI:** 10.1101/2021.04.08.438915

**Authors:** Laura Solé, Teresa Lobo-Jarne, Alberto Villanueva, Anna Vert, Yolanda Guillén, Irene Sangrador, Antonio Barbachano, Joan Lop, Marta Guix, Marta Salido, Beatriz Bellosillo, Raquel García-Romero, Marta Garrido, Jessica González, María Martínez-Iniesta, Erika Lopez-Arribillaga, Ramón Salazar, Clara Montagut, Ferrán Torres, Mar Iglesias, Toni Celià-Terrassa, Alberto Muñoz, Anna Bigas, Lluís Espinosa

## Abstract

Current therapy against colorectal cancer is based on DNA-damaging agents that eradicate highly proliferative malignant cells. Whether sublethal chemotherapy affects tumor cell behavior and impacts on patient outcome is primarily unstudied. We now show that sublethal chemotherapy imposes a quiescent-like state to p53 wildtype human colorectal cancer (CRC) cells that is linked to the acquisition of a fetal phenotype downstream of YAP1, similar to that observed after intestinal damage. CRC cells displaying this fetal phenotype exhibit tumor-initiating activity comparable to untreated cells but superior metastatic capacity. Notably, nuclear YAP1 accumulation, or detection of the fetal signature in tumors predict poor prognosis in CRC patients carrying p53 wildtype tumors. Collectively, our results uncover a potential adverse response of tumor cells to suboptimal chemotherapy, and identify nuclear YAP1 and fetal conversion of colorectal tumors as biomarkers for prognosis and therapy prescription.

**Statement of significance:** Chemotherapy induces a quiescent-like phenotype to colorectal cancer cells that is linked to the acquisition of a YAP1-dependent fetal signature. Notably, this signature is predictive of patient outcome in different cohorts of human colorectal cancer.

## INTRODUCTION

Colorectal cancer (CRC) remains the second leading cause of cancer-related death, which highlights the need for novel therapies focused on treatment of advanced disease. Treatment of localized CRC currently involves surgery, radiotherapy and/or chemotherapy (CT) (mainly 5-FU or capecitabine and oxaliplatin in the neoadjuvant or adjuvant setting), while CT (5-FU, oxaliplatin and irinotecan) still represents the main backbone of treatment for advanced CRC. In general, classical CT agents are designed to eradicate tumors by inducing DNA damage in highly proliferative cells leading to cell death (reviewed in (1)). However, most tumors contain a variable proportion of quiescent cells, including cancer stem cells, that are refractory to these agents thus contributing to tumor relapse and metastasis (2). In agreement with this notion, the presence of intestinal stem cell (ISC) signatures in tumors is predictive of poor prognosis in patients (3). Even after adequate treatment, around 25-30% of CRC patients in the less aggressive stage II tumors and up to 30-50% in stage III relapse and most of them eventually die (data from the American Cancer Society). There is growing evidence that therapeutic strategies that potentiate the effect of DNA damaging agents may provide the base for more effective combination therapies (4, 5).

Even when therapy fails to totally eradicate tumors, cancer cells receiving sublethal doses of therapeutic agents can acquire a senescent phenotype, characterized by high levels of the cell cycle inhibitors p16 and p21, cessation of proliferation and presence of a senescent-associated-secretory-phenotype (SASP) that effectively delays disease progression (6). However, senescent cells may still contribute to tumor progression as a source for relapse or metastasis, or by the secretion of pro-tumorigenic factors (reviewed in (7–9)). To date, there is an almost absolute lack of patient-based data to firmly establish the contribution of sub-lethal chemotherapy to patient outcome.

Recently, it was shown that fetal reprograming of intestinal cancer cells induced by YAP1 led to tumor and metastasis suppression in the Apc-/-; KrasG12D; p53-/- murine cancer model (10). We here show that p53 wildtype CRC patient-derived organoids (PDO)s treated with low doses of CT acquire a quiescent-like phenotype that is associated with YAP1-dependent fetal ISC conversion. These persistent quiescent-like (PQL) cells display high in vitro and in vivo tumor and metastasis initiating capacity, and we identified a restricted fetal ISC signature that is present in PQL cells and in a subset of untreated CRC tumors. Presence of this fetal signature or detection of nuclear YAP1 in tumors predicts poor disease outcome at stages II and III, in particular in CRC patients carrying tumors with functional p53 signaling.

## RESULTS

### Low-dose CT treatment of colorectal cancer PDOs induces a non-senescent quiescent phenotype in the absence of sustained DNA damage

To investigate the mechanisms that impose therapy resistant in cancer patients, we treated CRC PDOs with serial dilutions of the first-line CT agents 5-FU+Iri. Although high 5-FU+Iri. concentrations led to eradication of most PDO cells, we defined IC_20_ and IC_30_ as the 5-FU+Iri. doses that reduced cell viability by 20 and 30% after 72 hours of treatment, which were specific for each PDO (Figure 1A and Supplementary Table S1). Microscopy analysis of PDO5 (*TP53* WT) treated at IC_20_ and IC_30_ did not reveal obvious signs of cell death, but we noticed a dose-dependent growth arrest in all PDOs tested, that continued for at least 2 weeks after drug washout (Figures 1B and C, and S1A). Growth arrest was associated with inhibition of cell proliferation as determined by IHC analysis of ki67 (Figure 1D and S1B) and the reduced number of cells in S phase with accumulation in G_0_/G_1_ and G_2_/M (Figure S1C), the latter probably corresponding to cells not undergoing cytokinesis (11, 12). By fluorescent in-<u>s</u>itu <u>h</u>ybridization (FISH) and DAPI staining combined with IF of the membrane marker EPHB2, we demonstrated the absence of polyploid or multinucleated cells, respectively, following IC_30_ treatment (Figure S1D, E, F).

**Figure 1.**
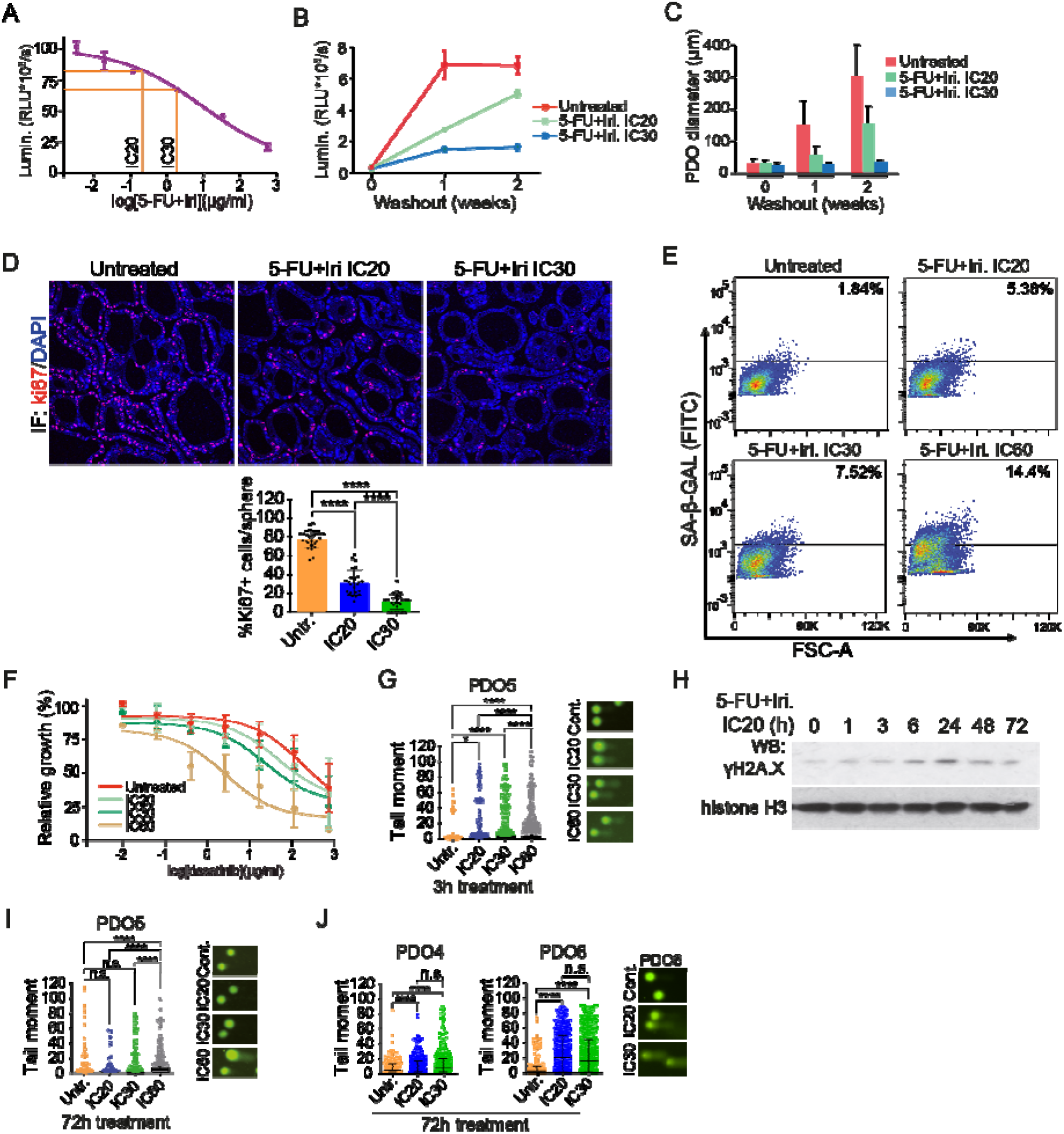
Low-dose CT treatment induces a non-senescent quiescent-like state to CRC PDO in the absence of persistent DNA damage. **(A)** Dose-response assay of PDO5 treated with 5-FU+Iri. for 72 hours, indicating the IC_20_ and IC_30_ doses. **(B and C)** Quantification of PDO5 (B) viability and (C) diameter, after 72 hours of 5-FU+Iri. treatment and subsequent washout and culture in fresh medium for 1 and 2 weeks. Representative data from 4 independent experiments is shown. **(D)** Representative images (upper panel) of ki67 staining by immunofluorescence (IF) in PDO5 tumoroid treated with 5-FU+Iri. at IC_20_ and IC_30_ for 72 hours, and quantification (lower panel) of the percentage of ki67^+^ cells/sphere in each condition. Counterstained with DAPI. **(E)** Quantification of SA-β-Gal activity detected by flow cytometry in PDO5 cells treated as in **(D)**. Representative data from 2 independent experiments is shown. (F) Dose-response curves of PDO5 treated with the senolytic drug dasatinib for 3 days after pre-treatment with 5-FU+Iri. at the indicated doses for 72 hours. **(G)** Comet assay to measure levels of DNA damage in PDO5 treated for 3 hours as indicated. (H) Western blot (WB) analysis of the DNA damage sensor γH2A.X in control and 5-FU+Iri.- treated PDO5 cells collected at the indicated time points after treatment. (I and J) Comet assay to measure levels of DNA damage in **(I)** 53 WT PDO5 and (J) p53 mutants PDO4 and PDO8, treated for 72 hours as indicated. The statistical analysis in **(D)**, **(G)**, **(I)** and **(J)** was performed by one-way ANOVA test, comparing treated with untreated condition and treated conditions with each other. p values are indicated as *p<0.05 and ****p < 0.0001. n.s., no significant; 5-FU, 5-fluorouracil; Iri, irinotecan; SA-β-Gal, SA-β-Galactosidase; CT, control; IC_20_, IC_30_ and IC_60_, 5-FU+Iri. treatment that results in 20%, 30% and 60% cell death, respectively, compared with untreated cell growth.

We determined whether IC_20_ and IC_30_ treatments inflicted a senescent phenotype to PDO5 cells by evaluation of <u>s</u>enescence-<u>a</u>ssociated (SA)-β-Galactosidase activity by flow cytometry (Figure 1E) and IHC (Figure S1G). Cells that persisted after IC_20_ or IC_30_ were not senescent, in contrast with cells treated at IC_60_ for 72 hours (Figure 1E). Accordingly, addition of the senolytic agent dasatinib (13) did not potentiate the growth inhibition imposed by IC_20_ and IC_30_ 5-FU+Iri. but enhanced the effect of IC_60_ 5-FU+Iri. treatment (Figure 1F). Moreover, we did not detect apoptotic PDO5 cells after IC_30_ treatment as determined by cleaved-caspase 3 staining (Figure S1H) and Annexin V staining (Figure S1I). We studied the possibility that cell cycle arrest after IC_20_ and IC_30_ treatment was linked to sustained DNA damage. Comet assay (Figures 1G) and WB analysis of the γH2A.X marker (Figure 1H) in PDO5 revealed a dose-dependent accumulation of DNA damage starting at 1-3 hours with a maximum at 24 hours. Importantly, DNA damage was undetectable at 72 hours after IC_20_ and IC_30_ treatment, but clearly present in IC_60_-treated PDO5 (Figure 1H and 1I). In contrast, PDO4 and PDO8 cells carrying mutated *TP53* exhibited high amounts of DNA damage following IC_20_ and IC_30_ 5-FU+Iri. treatment that lasted for at least 72 hours (Figure 1J), in agreement with the higher growth inhibition of *TP53* mutant PDOs after CT washout (Figure S1A).

These results indicate that p53 wildtype cancer cells that persisted after low CT acquire a quiescent-like phenotype, hereafter referred as PQL (for persistent quiescent-like), in the absence of sustained DNA damage.

### CT-induced PQL cells display fetal intestinal stem cells (feISC) characteristics

Sublethal CT treatment has been linked to the acquisition of specific stem cell signatures in B-cell lymphoma (8) and intestinal cancer (14). To study the transcriptional changes associated with the PQL phenotype, we performed RNA sequencing (RNA-seq) of control, IC_20_- and IC_30_- treated PDO5 cells. Bioinformatic examination of differentially expressed genes (DEGs) (Supplementary Table S2) showed an almost perfect correlation of gene expression changes between pairwise comparisons (IC_20_ vs. untreated and IC_30_ vs. untreated) (p<2.2e-16, R=0.974) (Figure S2A). Gene Set Enrichment Analysis (GSEA) uncovered p53 as the main activated pathway in PQL cells (Figure 2A), which was confirmed by WB analysis (Figure 2B), qPCR (Figure 2C) and ChIP assay (Figure S2B) of canonical p53 targets. DEGs genes also clustered in the NF-κB, epithelial-to-mesenchymal transition (EMT) and interferon gamma (IFNγ) pathways (Figure 2A) that have been associated with inflammatory response and stemness (15–19). Unexpectedly, DEG in our analysis negatively correlated with a canonical ISC signature (Muñoz et al., 2012) (Figure 2D). More in-depth analysis showed a mixed pattern of genes up-regulated such as *LY6D* and *YAP1*, which are instrumental in the fetal ISC (feISC) after intestinal injury (10,19–21), and down-regulated such is the case of canonical adult ISC markers *LGR5* and *EPHB2* (Figure 2E). Accordingly, GSEA indicated a significant correlation between the CT-induced signature and the transcriptional program associated with fetal ISC conversion (21) (Figure 2F and 2G).

**Figure 2.**
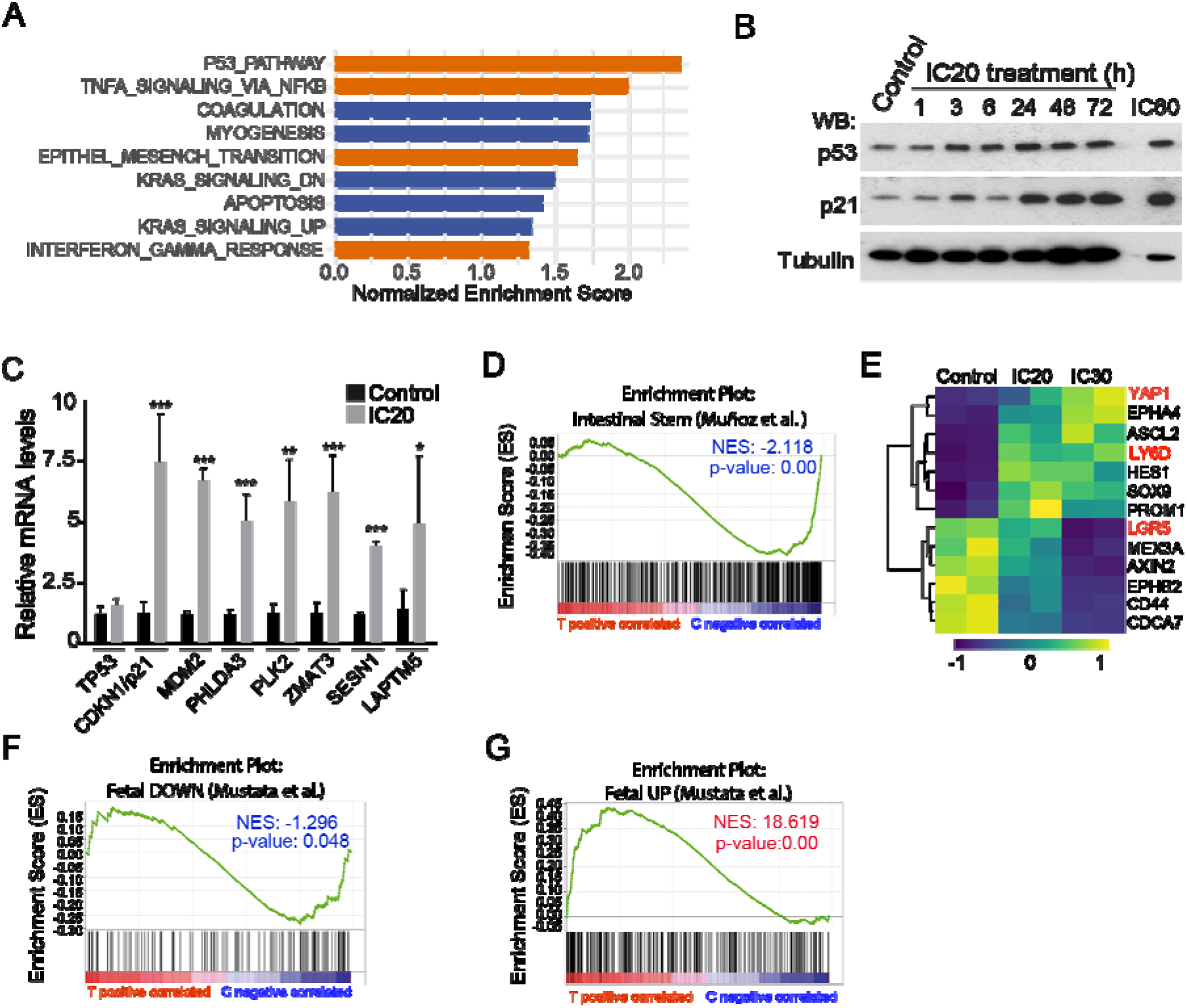
PQL phenotype is associated with acquisition of a fetal intestinal stem cell (feISC) signature. **(A)** Barplot depicting the normalized enrichment score of the statistically significant enriched pathways obtained by GSEA analysis with the Hallmark gene set for treated samples (NOM p- val<0.05). **(B)** WB analysis of control and treated PDO5 cells collected at the indicated time points after 5-FU+Iri. treatment. **(C)** RT-qPCR analysis of selected p53 target genes from control and IC_20_-treated PDO5 cells. **(D)** GSEA of an intestinal stem cell (ISC) gene set, according to Muñoz et al, in treated versus IC_20_-treated PDO5 condition. **(E)** Heat map showing the expression levels of the indicated ISC genes in untreated, IC_20_ and IC_30_-treated PDO5 cells. **(F and G)** GSEA of **(F)** a fetal down and **(G)** a fetal up stem cell gene set, according to Mustata et al, in control (C) versus treated (T) PDO5 condition. The statistical analysis in (C) was performed by T-Student test, comparing treated with untreated condition. p values are indicated as *p<0.05, **p < 0.01 and ****p < 0.0001. n.s., no significant; CT, control; IC_20_ and IC_30_, 5-FU+Iri. treatment that results in 20% and 30% cell death, respectively, compared with untreated cell growth; GSEA, gene set enrichment analysis; NES, normalized enriched score.

### The feISC signature shows a coordinate expression in human CRC and is dependent of functional p53

We asked whether CT-induced feISC signature was present in untreated human tumors. Computational analysis of the Marisa (22) (GSE39582), Jorissen (23) (GSE14333) and TCGA (TCGA Portal) CRC datasets using CANCERTOOL (24) indicated that many genes of the signature were distributed in clusters of coordinate expression in untreated tumors (with either positive or negative correlation) (Supplementary Table S3), which we integrated in a new cluster containing 28 plus 8 genes that were either upregulated (28up) or downregulated (8down) in CT-treated PDOs and fetal-converted ISCs (Figure S3A). This new 28up+8down-feISC gene signature was present in Marisa (Figure 3A), Jorissen and TCGA CRC cohorts (not depicted). We confirmed regulation of several genes of the 28up+8down-feISC signature by RT-qPCR (Figure 3B) and WB analysis (Figure 3C) of IC_20_ 5-FU+Iri.-treated PDO5. Activation of fetal genes following CT treatment was comparably observed in the *TP53* WT PDO66 (Figure 3D) but significantly impaired in the hypomorphic *TP53* mutant PDO4 (Figure 3E). We confirmed the p53 dependency of the feISC signature through generation by CRISPR-Cas9 of PDO5 pools with variable degrees of p53 KO (Figure S3B). RT-qPCR analysis revealed that PDO5 KO#3 with the highest p53 KO efficiency showed lower activation of 28up-feISC genes after 5-FU+iri. IC_20_ treatment (Figure 3F). However, we only detected slight differences in the protein levels of randomly selected fetal markers when comparing CRC cell lines carrying WT or mutant p53, with the latter showing a massive accumulation of the DNA damage marker γH2A.X after CT (Figure 3G). ChIP-seq assay of 5-FU+Iri. IC_20_-treated PDO5 cells indicated that only 3 genes in the 28up-feISC signature, PLK2, PHLDA3 and GSN genes were direct p53 targets (Figure S3C), consistent with previously published data (25). These results suggested that additional transcription factor/s govern fetal ISC conversion by CT.

**Figure 3.**
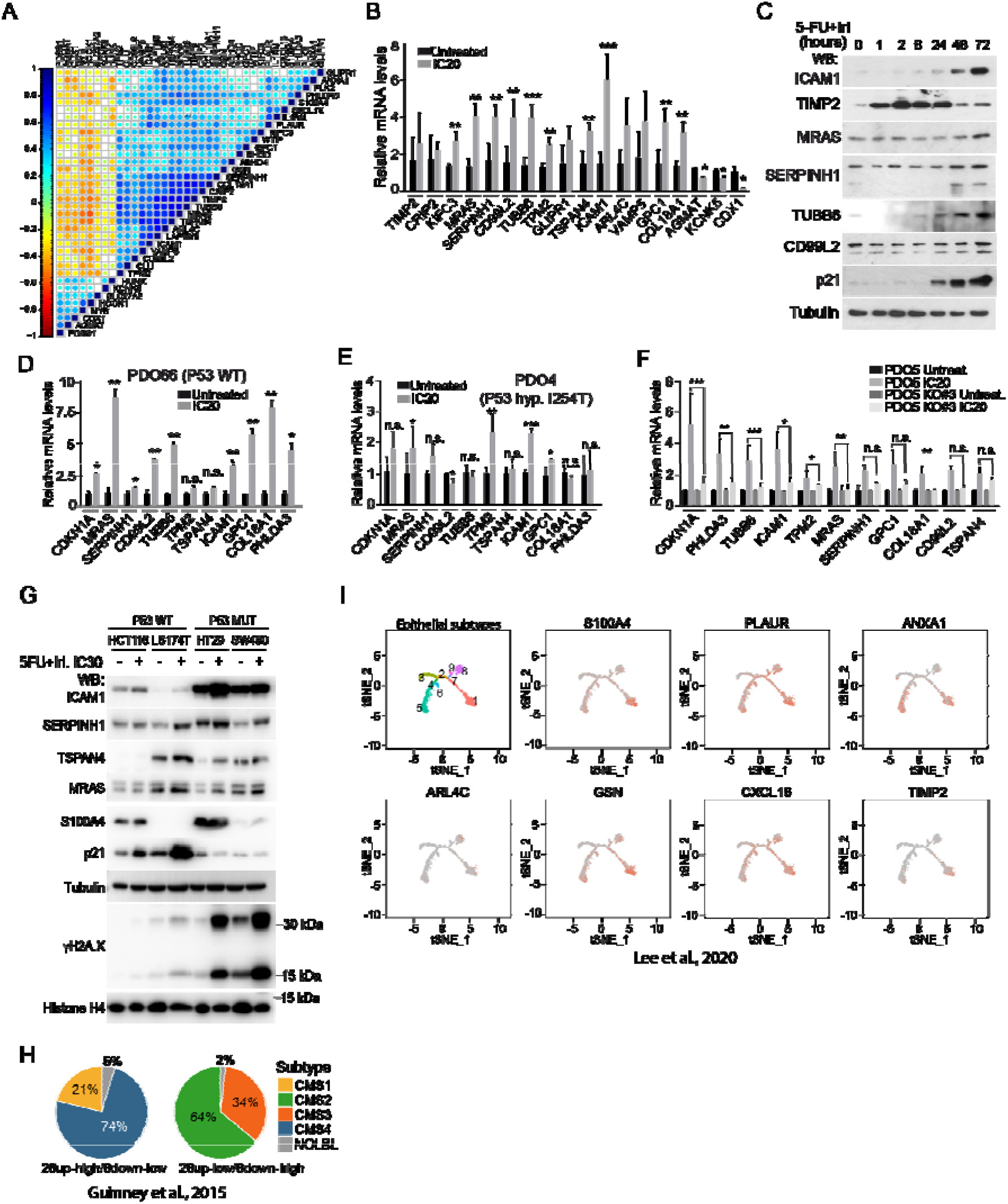
CT-induced quiescent cells display a fetal intestinal stem cell signature that is p53 dependent. **(A)** Expression correlation matrix from the 28up+8down-feISC gene signature using the Marisa database (*n=*566). The size of circles and color intensity are proportional to the Pearson correlation coefficient found for each gene pair. **(B)** RT-qPCR analysis of normalized relative expression of selected 28up+8down-feISC signature genes in control and treated PDO5 as indicated. **(C)** WB analysis of the indicated antibodies in control and treated PDO5 cells collected at the indicated time points after 5-FU+Iri. treatment. **(D, E and F)** RT-qPCR analysis of normalized relative expression of selected 28up-feISC signature genes plus CDKN1A gene in **(D)** control and CT-treated *TP53* WT PDO66, **(E)** *TP53* mutant PDO4 and **(F)** PDO5 *TP53* KO #3. **(G)** WB analysis of various CRC cell lines untreated or collected after 72 hours of 5-FU+Iri. treatment. Membranes were incubated with the indicated antibodies. **(H)** Pie charts showing the molecular subtype distribution (percentage), according to Guinney et al, in patients within the feISC signature groups as indicated. **(I)** Localization of several 28up-fetal-ISC genes in epithelial subtypes cell states 1-9 previously classified in Lee et al 2020. The *t*-SNE plots were obtained using the web-based tool URECA (User-friendly InteRface tool to Explore Cell Atlas). Cell states 1, 5 and 6 correspond to a transcriptional group enriched for secretory and migratory gene expression, whereas cell states 2, 3, 4, 7, 8 and 9 correspond to transport and Wnt signaling gene expression (27). The statistical analysis in (B), (D), (E) and (F) was performed by T-Student test, comparing treated with untreated condition. p values are indicated as *p<0.05, **p < 0.01 and ****p < 0.0001. n.s., no significant. CT, control; IC_20_ and IC_30_, 5-FU+Iri. treatment that results in 20% and 30% cell death, respectively, compared with untreated cell growth.

We also determined whether untreated tumors carrying the 28up+8down-feISC signature were restricted to a specific molecular cancer subtype. 74% of tumors with the 28up+8down-feISC signature were categorized as CSM4, based on the classification by Guinney and collaborators (26) (Figure 3H), which is characterized by upregulation of epithelial to mesenchymal transition gene signatures, TGFβ signaling, stromal infiltration and poorer patient prognosis. In contrast, 28up-low+8down-high tumors were primarily ascribed to the more canonical Wnt and Myc-driven CMS2 subtype. We studied whether the feISC signature of untreated tumors was expressed in the epithelial cancer cells or primarily contributed by the stromal component. Analysis of single cell RNA-seq data from Lee and collaborators (27) demonstrated that feISC genes are expressed in the epithelial cancer cells, particularly in states 1, 5 and 6 that are all associated with the secretory and migratory pathways (Figure 3I).

These results indicate that sublethal CT induces a fetal signature, which is dependent on the presence of a functional p53 pathway. This feISC signature is also expressed in a coordinate manner in untreated human CRC tumors, in particular in tumors of the CMS4 subtype from Guinney and collaborators, and the secretory and migratory epithelial states 1, 5 and 6 from Lee and collaborators.

### PQL cancer cells display increased in vitro and in vivo tumor initiation capacity

It was recently shown that fetal ISC conversion imposed by YAP1 activation (induced by deletion of the Hippo kinases LATS1/2) in the Apc^-/-^; Kras^G12D^; p53^-/-^ murine intestinal tumor cells results in tumor and metastasis suppression (10). Thus, we studied whether p53 proficient PQL cells, which display a consistent activation of a feISC signature, preserved tumor initiation capacity (TIC) of untreated cancer cells. We seeded 300 single cells from untreated or 5-FU+Iri.

IC_20_ or IC_30_-treated PDO5, as indicated. We found that CT pretreatment of PDO5 cells did not affect their TIC in vitro compared with untreated cells, as indicated by the equivalent number of spheres generated (Figures 4A, upper panel), but imposed a dose-dependent reduction of spheres diameter (Figures 4A, lower panel), consistent with their low proliferation rates. In contrast, 5- FU+Iri. pretreatment (IC_20_) of *TP53* mutant PDO4 and PDO8 cells resulted in TIC abrogation (Figure 4B), which is in agreement with the massive accumulation of DNA damage detected in comet assays (see Figure 1J). Considering that TIC activity in PDO5 could be driven by the fraction of cells which still undergo replication after IC_20_ and IC_30_ treatment (Figure 1D, S1B and S1C), we next compared the TIC in vitro of the general population and specifically the quiescent cells. For this, we generated *TP53* WT PDO5 carrying a doxycycline-inducible histone-GFP reporter that has been previously demonstrated to label the quiescent tumor population after doxycycline withdrawal (28). Upon 6 days of doxycycline treatment, PDO5 cells were treated with 5-FU+Iri. for 72 hours and, after 2 weeks of doxycycline washout, analyzed by flow cytometry and GFP_high_ and GFP_low_ were sorted (S4A). We found that sorted GFP_high_, which represents the quiescent population of CT-treated cells, displayed identical capacity for organoid generation as GFP_high_ plus GFP_low_ cells indicating that TIC activity is retained in the PQL population.

**Figure 4.**
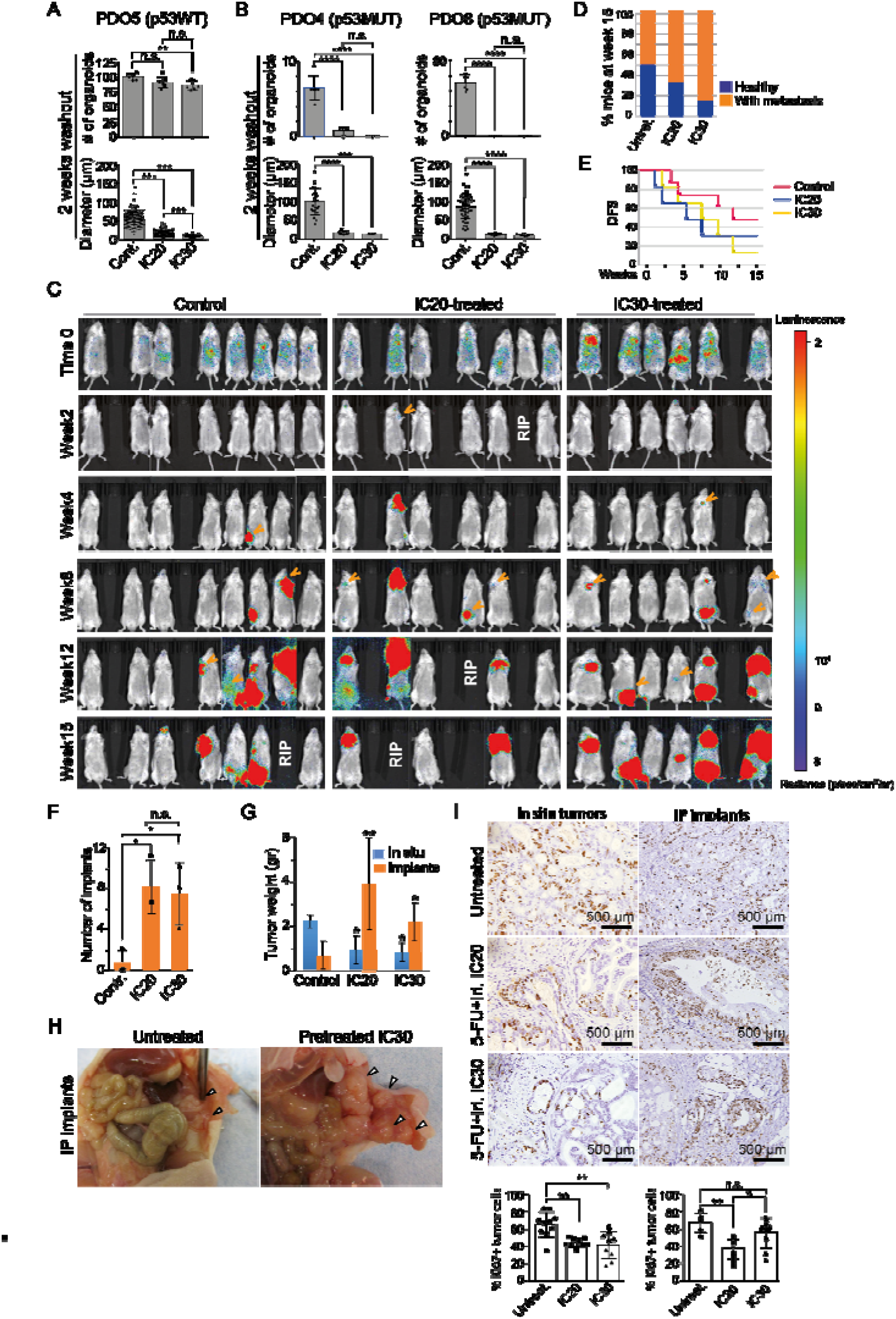
*TP53* WT PQL cells retain tumor-initiating capacity in vitro and in vivo. (A and B) Number of PDOs (upper panels) and diameter measurement (lower panels) of (**A)** *TP53* WT PDO5 and **(B)** *TP53* mutant PDO4 and PDO8 treated with 5-FU+Iri. as indicated and seeded at 300 cells/well as single cells and after 2 weeks with fresh medium. **(C)** In vivo bioluminescence representative images of mice administered intracardiac injection of 40.000 luciferase-PDO5 CT and IC_20_ or IC_30_ -treated cells in NSG mice. **(D)** Percentage of healthy and with metastasis mice at week 15 from (C). **(E)** Kaplan-Meier representation of disease-free survival probability over time for untreated, IC_20_ and IC_30_-treated PDO inoculated intracardiac in mice. **(F, G and H) (F)** Number of intraperitoneal implants, **(G)** tumor weight of in situ growing tumors and intraperitoneal implants and **(H)** photographs of tumors derived from orthotopically implanted CT, IC_20_ and IC_30_-pretreated PDOs in nude mice. **(I)** Immunohistochemistry (IHC) analysis of Ki67 in PDO-derived in situ tumors and implants and quantification of the percentage of Ki67^+^ tumor cells in the indicated conditions. The statistical analysis in (A), (B) and (I) was performed by one-way ANOVA test comparing treated with untreated conditions and treated conditions with each other. The statistical analysis in (F) and (G) by T-Student test, comparing treated with untreated condition. p values are indicated as *p<0.05 and **p < 0.01. n.s., no significant; 5-FU, 5-fluorouracil; Iri, irinotecan; DFS, disease-free survival; IC_20_ and IC_30_, 5-FU+Iri. treatment that results in 20% and 30% growth reduction, respectively, compared with untreated cell growth.

We next studied the in vivo tumorigenic capacity of IC_20_ and IC_30_ pretreated PDO5 cells using two complementary strategies. Firstly, we performed intracardiac injection of 40.000 single PDO5 cells (untreated, IC_20_ or IC_30_ pretreated) labelled with firefly luciferase into NOD-SCID-gamma (NSG) immunocompromised mice. Mice were analyzed weekly using bioluminescence (BLI) to monitor metastatic growth using the IVIS animal imaging system (Figure 4C). We found that PDO5 treated with 5-FU+Iri. displayed a slightly superior metastatic capacity than untreated cells. Specifically, 4 of 8 mice transplanted with untreated PDO5 cells contained metastatic lesions at week 15 after transplantation. Importantly, 4 of 6 mice transplanted with IC_20_-treated cells and 5 of 6 mice with IC_30_-treated cells showed visible metastasis 15 weeks after injection (Figure 4D and 4E). Secondly, we inoculated equivalent numbers of untreated, IC_20_ and IC_30_ pretreated PDO5 cells in the cecum of nude mice. Tumor growth was assessed by palpation weekly and animals sacrificed synchronously 70 days after transplantation. We found that untreated, IC_20_ and IC_30_-treated PDOs all generated tumors in the site of inoculation, being IC_20_ and IC_30_-treated derived tumors being much smaller than those arising from untreated controls (Figure 4F), as expected. Importantly IC_20_ and IC_30_-treated PDO cells displayed a significantly higher ability to generate intraperitoneal implants when compared with untreated tumor cells (Figure 4G and 4H). Still, we detected a reduction in the proliferation capacity of CT-treated PDO5 cells as determined by IHC analysis of the proliferation marker ki67 (Figure 4I).

These results indicate that *TP53* WT CRC cells treated with low doses of 5-FU+Iri. show reduced capacity to proliferate in vitro and in the primary tumors, but display comparable TIC as untreated cells in vitro and higher metastatic activity in vivo.

### The CT induced feISC signature is predictive of reduced disease-free survival in *TP53* WT tumors

We studied the possibility that the presence of the feISC signature in untreated CRC tumors was associated with patients’ outcome. To this aim, we analyzed the predictive capacity of the 28up+8down-feISC gene signature in the Marisa, Jorissen and TCGA CRC data sets. The global 28up+8down-feISC signature was sufficient to demarcate at least 2 subsets of patients in either data set (Figure 5A and Supplementary Table S4), with the group with highest 28up and lowest 8down-feISC levels displaying the poorest disease-free-survival (Figure 5B). A more detailed analysis of the Marisa data set demonstrated that this signature was significantly associated with tumor relapse in patients at stages II (n=264) (p=0.041) (Figure 5C) and II+III (n=469) (p=0.0033) (Figure 5C and 5D), and imposed a trend towards poor prognosis at stage IV (n=60) (Figure 5E).

**Figure 5.**
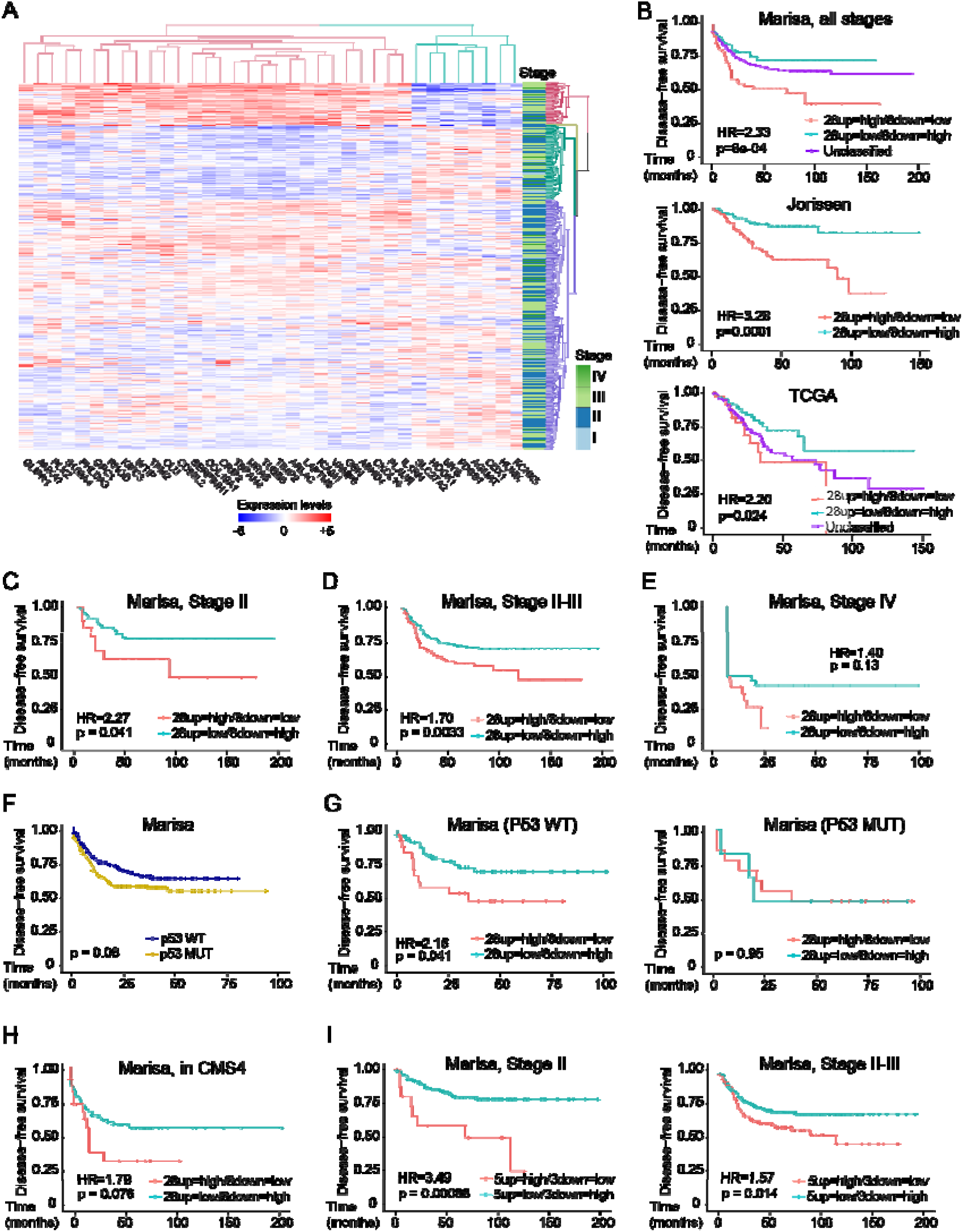
Identification of a fetal ISC signature with prognosis value in CRC. **(A)** Cluster analysis of the 28up+8down-feISC signature used to classify patients into at least two subsets in the Marisa colorectal cancer database. We allowed unsupervised hierarchical clustering of the 28+8 genes, while we enforced the classification of patients into 4 subsets (colored in red, green, light blue and purple). Positive and negative correlation expression levels is shown in red and blue, respectively. **(B)** Kaplan-Meier representation of disease-free survival probability over time for patients unclassified and with high or low expression of the 28up+8down-feISC signature selected according to (A) for Marisa colorectal cancer database (28up=high/8down=low *n=*66, 28up=low/8down=high *n=*114 and Unclassified *n=*385). Jorissen (28up=high/8down=low *n=*114 and 28up=low/8down=high *n=*112) and TCGA (28up=high/8down=low *n=*39, 28up=low/8down=high *n=*96 and Unclassified *n=*194) colorectal cancer databases were selected according to their cluster analysis of the 28up+8down-feISC signature. **(C, D and E)** Kaplan– Meier curves representing the disease-free survival probability over time for patients classified according to their cluster analysis of the 28up+8down-feISC signature of patient groups from **(C)** stage II (28up=high/8down=low *n=*23 and 28up=low/8down=high *n=*126), **(D)** stage II and III (28up=high/8down=low *n=*100 and 28up=low/8down=high *n=*368) and **(E)** stage IV (28up=high/8down=low *n=*25 and 28up=low/8down=high *n=*35), from Marisa colorectal cancer database. **(F)** Kaplan-Meier representation of disease-free survival probability over time of patents classified according to their *TP53* status in the Marisa colorectal cancer dataset (*TP53* WT n=161 and *TP53* MUT n=190). **(G)** Kaplan-Meier representation of disease-free survival probability over time of patients, from the Marisa dataset, classified according to their cluster analysis of the 28up+8down-feISC signature for patient groups from *TP53* WT (28up=high/8down=low *n=*27 and 28up=low/8down=high *n=*58) and *TP53* mutant (28up=high/8down=low *n=*6 and 28up=low/8down=high *n=*14). **(H)** Kaplan-Meier representation of disease-free survival probability over time of Marisa patient’s tumors previously categorized as CMS4 (26), and classified according to their cluster analysis of the 28up+8down-feISC signature (28up=high/8down=low *n=*23 and 28up=low/8down=high *n=*68). **(I)** Kaplan-Meier representation of disease-free survival probability over time for patients with high or low expression of the 5up+3down-feISC signature of stage III (5up=high/3down=low *n=*17 and 5up=low/3down=high *n=*156) and stage II and III (5up=high/3down=low *n=*87 and 5up=low/3down=high *n=*382) Marisa database, selected according to their unsupervised hierarchical cluster analysis. Data in (A) show normalized, centered and scaled Illumina probe set intensities on a log_2_ scale. The stage lane represents the stage subtype that corresponds to each patient. For statistical analysis of the Kaplan-Meier estimates we used Cox proportional hazards models (See Supplementary Table S4). HR, hazard ratio.; p, p-value. See also **Figure S5**.

Since presence of functional p53 defined feISC conversion and TIC maintenance (see Figures 3F, 4A and 4B), we explored the possibility that *TP53* status determined the prognosis value of 28up+8down-feISC signature in CRC patients. Stratification of Marisa (Figure 5F) and TCGA (Figure S5A) patients according to *TP53* status did not have prognosis value by itself, but determines the prognosis value of 28up+8down-feISC signature (Figures 5G and Figure S5B). Because 28up+8down-feISC tumors are mainly included in the worst prognosis CMS4, we tested whether this feISC signature represents an independent prognosis factor inside this molecular subtype. Our results indicate that the 28up+8down-feISC signature increased the risk of relapse in patients within the CMS4 group (Figure 5H).

Finally, we explored the possibility of identifying a simplified signature derived from the 28up+8down-feISC with comparable prognosis value in cancer, which would facilitate its implantation in the clinical practice. We scored genes by their coordinate expression in the 3 CRC datasets analyzed (Supplementary Table S5). Then we evaluated the added value of single genes to the simplest signature composed by the highest scored 28up plus the highest scored 8down-feISC gene. We uncovered a minimal 5up+3down-feISC signature that shared coordinate expression in tumors (Figure S5C) and stratified patients with poor prognosis in all tested CRC cohorts (Marisa, p=0.00003; Jorissen, p=0.0002; TCGA, p=0.037) (Figure S5D) including patients carrying tumors at stage II and II-III (Figure 5I).

### Acquisition of feISC by CT treatment is linked to and dependent on YAP1 activation

We investigated whether feISC conversion of CRC cells after CT treatment was YAP1 dependent, as previously found in mouse models (10, 20). WB analysis of PDO5 cells (Figure 6A) and various CRC cell lines (Figure 6B) showed increased YAP1 expression after 5-FU+Iri. treatment, that was restricted to cells carrying WT *TP53* (Figure 6B). We detected accumulation of nuclear (active) YAP1 in IC_20_ and IC_30_-derived PDO5 tumors at 2 months after implantation in mice (Figure 6C). Next, we studied whether YAP1 activity was required for transcriptional induction of feISC genes by CT. Incubation of PDO5 cells with the YAP1 inhibitor verteporfin precluded induction of all tested feISC genes following IC_20_ 5-FU+Iri. treatment (Figure 6D). Similarly, IC_20_ 5-FU+Iri. treatment of Ls174T CRC cells led to an increase in YAP, SERPINH1 and TSPAN4 protein levels, which was abrogated by verteporfin (Figure S6A). Importantly, verteporfin alone, but more effectively in combination with CT, promotes tumor cell death in both *TP53* WT and *TP53* mutant PDO cells (Figures 6E). In addition, we observed that IC_20_ and IC_30_ pre-treated PDO cells show increased resistance to subsequent CT treatment (Figure S6B) that was prevented by addition of verteporfin (Figure 6F) strongly suggesting that combination of CT plus YAP1 inhibitors may represent a suitable therapeutic strategy for eradicating CRC tumors showing fetal ISC conversion.

**Figure 6.**
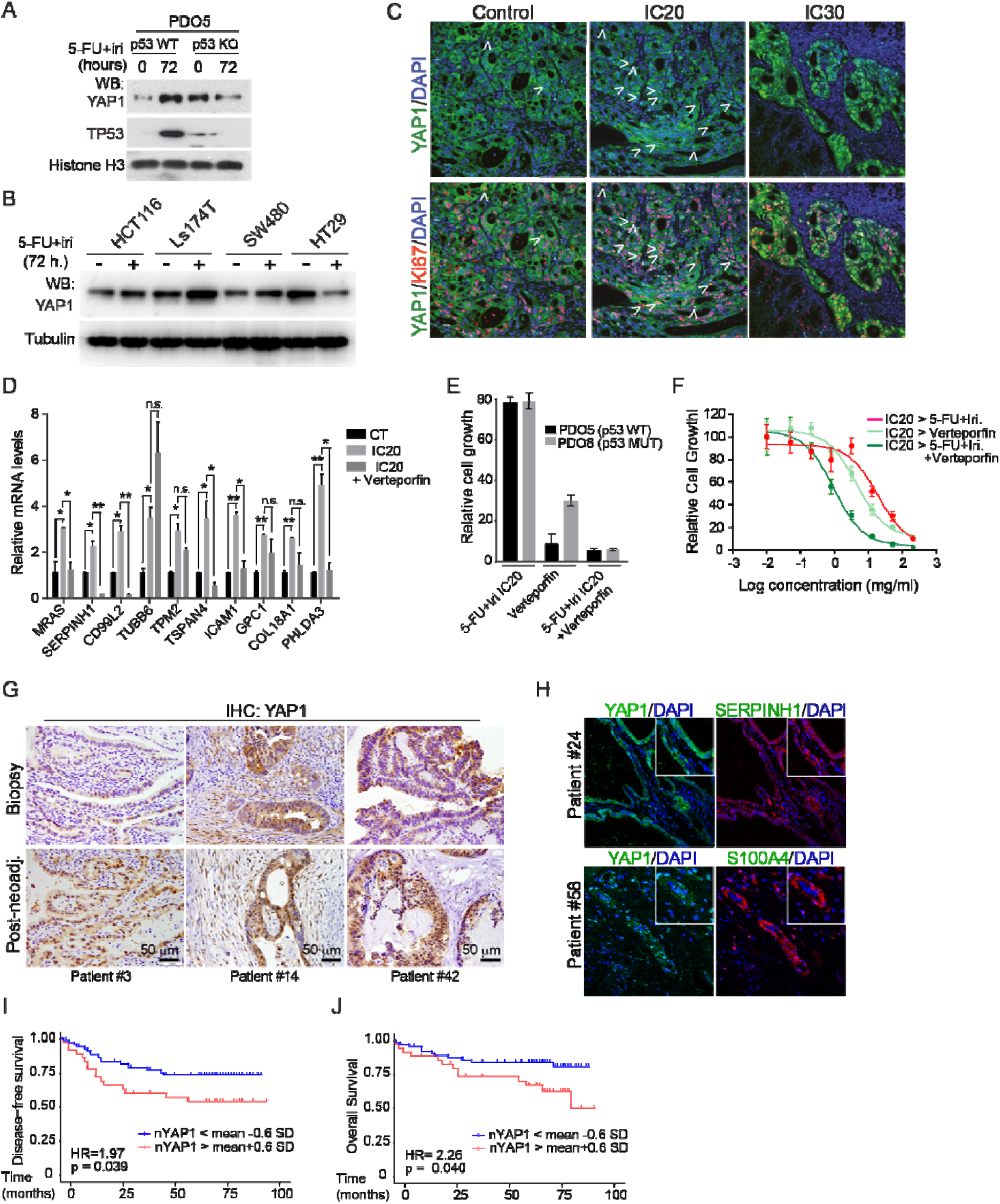
Acquisition of feISC by CT treatment is dependent on YAP1 activation. **(A)** WB analysis of the indicated antibodies of control and *TP53*-*depleted* PDO5 KO# 3 cells collected after 72 hours of 5-FU+Iri. treatment. **(B)** WB analysis of *TP53* WT (HCT116 and Ls174T) and *TP53* mutant (SW480 and HT29-M6) CRC cell lines untreated and collected after 72 hours of 5-FU+iri. treatment. **(C)** Representative images of YAP1 and the proliferation marker ki67 staining by IF in tumors derived from orthotopically implanted CT, IC_20_ and IC_30_- pretreated PDOs in nude mice. White arrows indicate nuclear translocation of YAP1. Counterstained, DAPI. **(D)** IHC analysis of YAP1 in representative type 1 (#14) and type 2 (#3, #42) colorectal tumor samples from the same patient at diagnosis (biopsy) and after neoadjuvant therapy at the time of surgery (post-neoadjuvancy). We classified patients based on ki67 levels in the tumor at diagnosis and at surgery. Complete results are shown in Supplementary Table S6. **(E)** Representative images of YAP1 and the fetal gene S100A4 by IF in type 1 (#24) and type 2 (#58) colorectal tumor samples (post-neoadjuvancy). **(F)** RT-qPCR analysis of normalized relative expression of selected 28up-feISC signature genes in control and treated PDO5 with 5- FU+Iri. alone or in combination with verteporfin at a final concentration of 0.2 μM. **(G)** Representative stereoscopic images (left panel) of the *TP53* WT PDO5 and *TP53* mutant PDO8 treated with 5-FU+Iri. alone, verteporfin alone or in combination with verteporfin at a final concentration of 1 μM and quantification (right panel) of the relative cell growth in both PDOs treated with verteporfin alone or in combination with 5-FU+Iri. **(H)** Dose-response curves of IC_20_-pretreated PDO5 treated for 3 days as indicated. Statistical analysis in (F) was performed by T-Student test, comparing treated with untreated conditions and treated conditions with each other. p values are indicated as *p<0.05, **p < 0.01, n.s., no significant. CT, control; IC_20_ and IC_30_, 5-FU+Iri. treatment that results in 20% and 30% cell death, respectively, compared with untreated cell growth.

Then, we performed IHC analysis of ki67 (to determine the proliferation status) and YAP1 in 62- paired human CRC tumors collected at diagnosis (biopsy) and after DNA damaging-based neoadjuvant treatment (surgery). Whereas some tumors exhibited similar proliferation rates after treatment, as determined by Ki67 staining (type 1), we identified a large subset of tumors that displayed reduced proliferation with no morphological evidences of senescence (type 2), such as enlarged nuclei or expression of the senescence marker p16 (Figure S6C), which are observed in scarce tumors at surgery (type 3) (Supplementary Table S6). We failed to detect differences in patient prognosis when comparing type 1 and type 2 tumors that are readily observed in patient carrying type 3 tumors (no events of relapse in the follow-up period) (Figure S6D). Interestingly, we detected nuclear YAP1 in few epithelial cells of untreated tumors, which was massively increased in neoadjuvant treated tumors independently of the proliferation status (Figure 6G and Supplementary Table S6), associated with expression of the feISC markers S100A4 and SERPINH1 (Figure 6H).

Since the number of samples in the studied cohort was insufficient to evaluate the clinical impact of nuclear YAP1 accumulation, we performed IHC analysis of YAP1 in a tissue microarray containing 196 different human CRC samples in triplicates with available clinical data. We determined the H score of nuclear YAP1 (as intensity multiplied by percent of positive tumor cells) in the triplicates and stratified the CRC patients accordingly. Considering the mean value ±

0.2 standard deviations of the H-score, we observed a trend towards poor prognosis in the group with higher nuclear YAP1 (disease-free survival: p=0.26; HR=1.38, not depicted) that increased when considering mean value ± 0.4 s.d. (p=0.12; HR=1.58, not depicted). Importantly, nuclear YAP1 levels reached statistical significance when considering the mean value ± 0.6 standard deviations (p=0.039; HR=1.97) (Figure 6I and 6J).

Together our data indicate that sublethal CT promotes the conversion of tumor cells into a fetal ISC phenotype that favors cancer progression and metastasis. Moreover, we uncover a CT-induced 28up+8down-feISC signature and its derivative the 5up+3down-feISC signature, which are present in a subset of untreated CRC tumors and has prognosis value in the context of functional p53. Finally, we demonstrated the higher efficacy of CT in combination with YAP1 inhibitors for eradication of *TP53* WT tumors, and the possibility of using nuclear YAP1 detection to identify patients that could benefit from this therapeutic strategy.

## DISCUSSION

CT is the current treatment for advanced and metastatic colorectal tumors. However, in a percent of cases tumor cells that escape from death imposed by therapeutic drugs (by efficient drug clearance, effective DNA repair or due to reduced accessibility of the drugs to specific tumor areas) can acquire a dormant phenotype that provides superior resistance to subsequent DNA damaging-based treatment based on their less proliferative state. In the present study we have shown that sublethal doses of CT impose a non-senescent and non-proliferating phenotype to cancer cells, in the absence of persistent DNA damage. Although discriminating between senescence and quiescence is not essential, since senescence was demonstrated to be reversible depending on the alternative use of p53/p21 (reversible) or p16/RB (irreversible) (29) leading to increased tumor stemness (8), we unequivocally observed that CT-treated cells display a quiescence-like state in the absence of a robust senescence phenotype. Importantly, PQL cancer cells can efficiently escape from dormancy following in vivo transplantation, as it is shown in the xenograft experiments, displaying a superior capacity to escape from the site of implantation. This is in agreement with the higher metastatic potential observed for CT treated cells and in line with previously studies showing that dormant cell populations in primary human CRC cells still retains tumor propagation potential (30) and that specific subpopulations of cancer cells reversibly enter a quiescent state and exhibit increased tumorigenic potential in response to CT (31).

Our transcriptomic studies revealed that the dynamics associated with the acquisition of the PQL phenotype rely downstream of p53 and p21 signaling. It has long been established that the key regulatory proteins that mediate cell cycle block include p53, p21 and p16, among others (reviewed in (32). The finding that the cancer cells carrying dysfunctional p53 did not acquire the PQL phenotype but display massive amounts of DNA damage may explain the results obtained by Cheung and collaborators indicating that YAP1 activation acts as tumor suppressor in p53 depleted tumors (10). In this context, our results are of particular relevance since they clarify the functional contribution of YAP1 as driver of fetal conversion and metastasis in p53 WT colorectal cancer, which is in agreement with previous reports (reviewed in (33)). In the same direction, it has been recently demonstrated that tumor cells that resist prolong CT acquire an embryonic-like and quiescent state that facilitates therapeutic resistance (34). We propose that functional p53 through p21 upregulation imposes a stop in proliferation when exposed to low CT doses that allow cells to recover under specific conditions, whereas cell depleted of functional p53 continue proliferating thus accumulating irreparable damage and apoptotic death. This cellular response represents, in fact, a double-edge sword since it can impose specific outcomes depending on the *TP53* status and the heterogeneity of cancer cells. In this sense, it was demonstrated that 5-FU treatment induced cell dormancy and epithelial to mesenchyme transition in lung cancer cells, associated with p53 accumulation (35). Further experiments genetically deleting YAP1 in *TP53* WT cells and preclinical assays using YAP1 inhibitors are required to definitively demonstrate the possibility of using specific protocols to combat fetal- converted tumors.

Remarkably, we have shown that the sublethal CT-induced feISC signature is already present in a subset of untreated tumors at diagnosis in several CRC cohorts. It is tempting to speculate that extrinsic factors or non-cancer cells present in the tumor, such as inflammatory cells, may induce the upstream regulators of this signature (i.e., TGFβ signaling) thus leading to the acquisition of PQL traits in the absence of treatment. In agreement with this idea, tumors carrying the 28up+8down-feISC signature are primarily included in the CMS4 CRC subtype identified by Guinney and collaborators (26) and characterized by stromal infiltration and TGFβ signaling. We speculate that TGFβ or additional cytokines derived from the tumor stroma may impose a YAP1-dependent feISC signature, which is in agreement with the previous demonstration that TGFβ promotes YAP1 signaling by facilitating the degradation of the negative regulator of the pathway RASSF1A (36). To recognize patients with higher probability of recurrence among those of uncertain prognosis (stages II-III) by the analysis of a reduced feISC signature can be clinically relevant as it may suggest more aggressive treatments or to intensify patient follow-up. Additionally, targeting the upstream signals imposing PQL/feISC acquisition pharmacologically (i.e. YAP1 inhibitors) or using combination treatments that effectively eradicate quiescent tumor cell populations (i.e. CT plus inhibitors of the NHEJ repair pathway) appear as interesting therapeutic options.

On the other hand, we identified IFN signaling in the analysis of CT-induced genes associated with PQL acquisition, which is in agreement with the essential function of IFN pathway in the conversion of adult stem cells into a fetal ISC phenotype (19). Conversion of adult into fetal ISC had already been identified as part of the process of tissue regeneration after helminths infection (19) or in the Dextran Sulfate Sodium colitis model (20). Thus, our results reinforce the concept that tumor development is partially mimicking the tissue regeneration process.

Our findings linking fetal ISC conversion poor CRC prognosis are partially in contrast with data indicating the prognosis factor of the adult ISC signature, including Lgr5 (3). Nevertheless, it has been recently demonstrated that Lgr5 and other adult ISC markers are temporary lost from cells seeding metastases, and subsequently recovered (due to cellular plasticity) to allow metastasis establishment (37) and most data indicating the requirement for adult Lgr5+ ISC in metastasis seeding have been obtained on p53-deficient tumor cells (10, 38). Interestingly, Batlle’s group recently identified a different contribution of Lgr5+ cells in PDOs carrying mutated or WT *TP53* (higher in mutant *TP53*) (39) thus opening the possibility that dependance on adult Lgr5+ ISCs for cancer progression is linked to *TP53* deficiency, which should be further investigated. Independently on the mechanisms underlying conversion of adult into fetal ISCs (induced by CT or signals derived from the tumor stroma), we have here identified a restricted genetic signature that is present in a subset of tumors that are mostly included in the CMS4 (the cancer stem cell and poor prognosis subtype) but clearly differ from that of adult ISCs.

From a clinical perspective, uncovering genetic signatures that are predictive of recurrence in a group of patients with uncertain projection (stages II and III) will represent a powerful tool for diagnosis refinement. In this direction, we are currently setting up the protocols for early detection of PQL/feISC cells in stages II-III tumors at diagnosis. As mentioned, anticipating the presence of this adverse phenotype in tumors would allow exposing selected groups of patients to alternative therapeutic procedures that could be refined with the discovery of the mechanisms imposing fetal SC conversion in cancer.

## METHODS

### Study design

The goal of this study was to determine in patient-derived data that chemotherapy positively or negatively impact on colorectal cancer progression in case of incomplete remission. Study of sublethal doses of DNA-damaging agents was performed in several patient-derived organoids and human cell lines to demonstrate its broad effects. By taking advantage of public colorectal tumor databases, the expression profiling of genetic signatures was accomplished. The numbers of experiments, biological replicates, and sample sizes for each database are outlined in the figure legends.

### Reagents, antibodies and software

A table of the source of all reagents, antibodies, kits, cell lines, chemicals and software is included (Table S8).

### Animal studies

Fragments of human colorectal tumors obtained from Parc de Salut MAR Biobank (MARbiobank) with the informed consent of patients and following all recommendations of Hospital del Mar’ Ethics Committee, the Spanish regulations, and the Helsinki declaration’s Guide were transplanted and expanded in the cecum of nude mice as orthoxenografts. To perform tumor-initiating assays, two approaches were used. Firstly, intracardiac injection of 40.000 CT (n=8) and IC_20_ (n=7) or IC_30_ (n=6) -treated PDO5 cells carrying a luciferase reporter to NSG mice was performed. For checking that the injection was performed correctly, after injection animals were anesthetized and were given 100µl of substrate D-luciferin at 15 mg/ml by intraorbital injection. Bioluminescent imaging was performed placing the animals into the IVIS Lumina III In Vivo Imaging System (PerkinElmer). Images were recorded with an exposure time of 2 minutes and were taken every week. Quantification was done using Living Image® software (PerkinElmer). Secondly, equivalent amounts of disaggregated patient-derived organoids (PDOs), previously treated as indicated below, were implanted as orthoxenografts. Follow-up of the growing tumors was done by palpation and animals were sacrificed when controls developed tumors of around 2 cm of diameter. In all our procedures, animals were kept under pathogen-free conditions, and animal work was conducted according to the guidelines from the Animal Care Committee at the Generalitat de Catalunya. The Committee for Animal Experimentation at the Institute of Biomedical Research of Bellvitge (Barcelona) approved these studies.

### Patient-derived organoids and culture conditions

Samples from patients were kindly provided by MARBiobank and IdiPAZ Biobank, integrated in the Spanish Hospital Biobanks Network (RetBioH; www.redbiobancos.es). Informed consent was obtained from all participants and protocols were approved by institutional ethical committees. For patient-derived organoids (PDOs) generation, primary or xenografted human colorectal tumors were disaggregated in 1.5 mg/mL collagenase II and 20 μg/mL hyaluronidase after 40min of incubation at 37°C, filtered in 100 μm cell strainer and seeded in 50 μL Matrigel in 24-well plates, as previously described (40). After polymerization, 450 μL of complete medium was added (DMEM/F12 plus penicillin (100 U/mL) and streptomycin (100 μg/mL), 100 μg/mL Primocin, 1X N2 and B27, 10mM Nicotinamide; 1.25 mM N-Acetyl-L-cysteine, 100 ng/mL Noggin and 100 ng/mL R-spondin-1, 10 μM Y-27632, 10 nM PGE2, 3 μM SB202190, 0.5 μM A-8301, 50 ng/mL EGF and 10 nM Gastrin I). Tumor spheres were collected and digested with an adequate amount of trypsin to single cells and re-plated in culture. Cultures were maintained at 37°C, 5% CO_2_ and medium changed every week. PDOs were expanded by serial passaging and kept frozen in liquid Nitrogen for being used in subsequent experiments. Mutations identified in the PDOs are listed in Supplementary Table S1.

### Cell lines

CRC cell lines HCT116 and Ls174T (*KRAS* mutated and *TP53* WT), SW480 (*KRAS* and *TP53* mutated) and HT29 (B*RAF* and *TP53* mutated) were obtained from the American Type Culture Collection (ATCC, USA). All cells were grown in Dulbecco’s modified Eagle’s medium (Invitrogen) plus 10% fetal bovine serum (Biological Industries) and were maintained in a 5% CO_2_ incubator at 37°C. 5-FU+Iri. concentrations that reduced 30% of each cell growth were as follows: HCT116, 0.01 µg/mL 5-FU and 0.004 µg/mL Iri.; Ls174T, 0.025 µg/mL 5-FU and 0.01 µg/mL Iri.; SW480, 0.28 µg/mL 5-FU and 0.11 µg/mL Iri.; HT29, 0.33 µg/mL 5-FU and 0.13 µg/mL Iri.

### Human colorectal cell lines

Formalin-fixed, paraffin-embedded tissue blocks of gastrointestinal tumor samples, from patients at diagnosis and after neoadjuvant therapy at the time of surgery, were obtained from Parc de Salut Mar Biobank (MARBiobank, Barcelona). Samples were retrieved under informed consent and approval of the Tumor Bank Committees according to Spanish ethical regulations and the guidelines of the Declaration of Helsinki. Patient identity for pathological specimens remained anonymous in the context of this study. Patient data was collected (Supplementary Table S6). IHC analyses were performed as described below.

### Patient-derived organoids viability assays

600 single PDO cells were plated in 96-well plates in 10 μL Matrigel with 100 μL of complete medium. After 6 days in culture, growing PDOs were treated with combinations of 5-FU+Iri. for 72 hours at the concentrations that reduce a 20 and 30% of the cell growth (IC_20_ and IC_30_, respectively), which are specific for each PDO as described in Supplementary Table S1. After 72 hours of treatment, we changed to fresh medium and measured the cell viability after 3 days, 1 week and 2 weeks using the CellTiter-Glo 3D Cell Viability Assay following manufacturer’s instructions in an Orion II multiplate luminometer. Images were obtained with an Olympus BX61 microscope at the indicated time points and the diameter of at least 70 tumoroids per condition was determined using Adobe Photoshop. For dose-response curves, PDOs were plated in 96-well plates in Matrigel and after 6 days in culture were treated with combinations of 5-FU and Irinotecan. Following 72 hours of treatment, we changed to fresh medium and treated with increasing concentrations of either 5-FU+Iri., dasatinib, verteporfin or combinations for 72 hours at the indicated concentrations. Cell viability was determined as described.

### Cell cycle analysis

Cell cycle was determined by flow cytometry using the standard APC BrdU Flow Kit. Briefly, treated PDOs with combinations of 5-FU+Iri., as indicated, were stained with bromodeoxyuridine (BrdU) for 24 hours. Single cells were obtained and processed according to the manufacturer’s instructions, with DAPI staining for the DNA content. The cells were analyzed in the LSR II analyzer.

### Cell senescence assays

Cell senescence was identified by the presence of SA-β-galactosidase activity using two different approaches. On one hand, staining for SA-β-galactosidase activity in cultured cells was carried out using the Senescence β-Galactosidase Staining Kit. Briefly, PDOs were seeded in 24-well plates (3000 cells per well). After 6 days, PDOs were treated with combinations of 5-FU+Iri. for 72 hours and were subsequent stained with the β-Galactosidase Staining Solution for 2 hours, according to the manufacturer’s instructions. Sections embedded in paraffin were counterstained with Fast Red for nuclei visualization. Images were obtained with an Olympus BX61 microscope. On the other hand, SA-β-galactosidase activity was addressed by flow cytometry using the Cell Event Senescence Green Flow Cytometry Assay Kit following the manufacturer’s instructions, and analyzed in the LSR II analyzer.

### Cell lysis and Western Blot

Treated PDOs were lysed for 20 min on ice in 300 μL of PBS plus 0.5% Triton X-100, 1 mM EDTA, 100 mM NA-orthovanadate, 0.2 mM phenyl-methylsulfonyl fluoride, and complete protease and phosphatase inhibitor cocktails. Lysates were analysed by western blotting using standard SDS–polyacrylamide gel electrophoresis (SDS-PAGE) techniques. In brief, protein samples were boiled in Laemmli buffer, run in polyacrylamide gels, and transferred onto polyvinylidene difluoride (PVDF) membranes. The membranes were incubated with the appropriate primary antibodies overnight at 4°C, washed and incubated with specific secondary horseradish peroxidase–linked antibodies. Peroxidase activity was visualized using the enhanced chemiluminescence reagent and autoradiography films.

### RT-qPCR analysis

Total RNA from treated PDOs was extracted with the RNeasy Micro Kit, and cDNA was produced with the RT-First Strand cDNA Synthesis Kit. RT-qPCR was performed in LightCycler 480 system using SYBR Green I Master Kit. Samples were normalized to the mean of the housekeeping genes *TBP* and *HPRT1*. Primers used for RT-qPCR are listed in Supplementary Table S7.

### ChIP-sequencing analysis

IC20-treated PDO5 was subjected to ChIP as previously described (41). Briefly, formaldehyde crosslinked cell extracts were sonicated, and chromatin fractions were incubated for 16 h with anti-p53 [abcam ab 1101] antibody in RIPA buffer and then precipitated with protein A/G-sepharose [GE Healthcare, Refs. 17-0618-01 and 17-0780-01]. Crosslinkage was reversed, and 6–10 ng of precipitated chromatin was directly sequenced in the genomics facility of Parc de Recerca Biomèdica de Barcelona (PRBB) using Illumina® HiSeq platform. Raw single-end 50- bp sequences were filtered by quality (Q > 30) and length (length > 20 bp) with Trim Galore (42). Filtered sequences were aligned against the reference genome (hg38) with Bowtie2 (43). MACS2 software (44) was run first for each replicate using unique alignments (q-value < 0.1). Peak annotation was performed with ChIPseeker package (45) and peak visualization was done with Integrative Genomics Viewer (IGV). ChIP- sequencing data are deposited at the GEO database with accession number GSE164161.

### RNA-sequencing experiments and data analysis

Total RNA from untreated and treated PDOs was extracted using RNeasy Micro Kit. The RNA concentration and integrity were determined using Agilent Bioanalyzer [Agilent Technologies]. Libraries were prepared at the Genomics Unit of PRBB (Barcelona, Spain) using standard protocols, and cDNA was sequenced using Illumina HiSeq platform, obtaining ∼ 45-64 million 50-bp paired< end reads per sample. Adapter sequences were trimmed with Trim Galore. Sequences were filtered by quality (*Q* > 30) and length (> 20 bp). Filtered reads were mapped against the latest release of the human reference genome (hg38) using default parameters of TopHat (v.2.1.1) (46) and expressed transcripts were then assembled. High-quality alignments were fed to HTSeq (v.0.9.1) (47) to estimate the normalized counts of each expressed gene. Differentially expressed genes between different conditions were explored using DESeq2 R package (v.1.24.0) (48) and adjusted P-values for multiple comparisons were calculated applying the Benjamini-Hochberg correction (FDR) (see Supplementary Table S2). Plots were done in R. Expression heatmaps were generating using the heatmaply and pheatmap packages in R (49). Gene Set Enrichment Analysis (GSEA) was performed with described gene sets using gene set permutations (n = 1000) for the assessment of significance and signal-to-noise metric for ranking genes. RNA-sequencing data are deposited at the GEO database with accession number GSE155354.

### Signature definition

To generate the fetal intestinal stem cell signatures, we selected genes with log2 Fold Change (FC) TreatedvsControl > 0 and FetalvsAdult (21) > 0 in the case of the 28up-feISC and log^2^FC TreatedvsControl < 0 and FetalvsAdult < 0 in the case of the 8down-feISC. Next, we used the Marisa data set to performed expression correlation matrices for the selected expression gene pairs using the corrplot package (v.0.84). To obtain the simplified signature genes were scored by their coordinate expression taking into account the three CRCR datasets analyzed (see Supplementary Table S5). Then it was evaluated adding a value of single genes to the simplest signature composed by the highest scored 28up plus the highest scored 8down-feISC. The process ended when adding a gene did not improved the prognosis value. Correlations were considered as statistically significant when the Pearson correlation coefficient corresponded to a p value below 0.05. Clusters of genes were selected when the absolute value for the Pearson correlation coefficient was above 0.1. Correlations were considered as statistically significant when the Pearson correlation coefficient corresponded to a p value below 0.05. Clusters of genes were selected when the absolute value for the Pearson correlation coefficient was above 0.1.

### Quantification and Statistical analysis

Statistical parameters, including number of events quantified, standard deviation and statistical significance, are reported in the figures and in the figure legends. Statistical analysis has been performed using GraphPad Prism 6 software, and P < 0.05 is considered significant. Two-sided Student’s t-test was used to compare differences between two groups. Each experiment shown in the manuscript has been repeated at least twice. Combinations of 5-FU+Iri. treatment has been checked for an appropriate IC20 and IC30 effect in every experiment, by cell viability assay. Bioinformatic analyses were performed as indicated above.

## Acknowledgments

We want to thank the Bigas’ and Espinosa’s lab members for constructive discussions and suggestions and technical support. We thank our patients for their generosity and to MARbiobank and the IdiPAZ Biobank integrated in the Spanish Hospital Biobanks Network (RetBioH; www.redbiobancos.es).

## Author contributions

Conceptualization: AB, LE Biochemical assays, in vitro and in vivo experiments: AV, MG, MS, RGR, MMI, IS Experiments, investigation and evaluation of results: LS, TLJ, AVe, TCT, AM Big data analysis and statistical analysis: TLJ, YG, ELA, FT Clinicopathological characterization of human tumors: MG, RS, CM. Clinical advice: MG, RS, CM Writing – original drat: LE, AB Writing -review & editing: all authors

## Data and materials availability

All data associated with this study are present in the paper or the Supplementary Materials. RNA sequencing and ChIP sequencing data have been deposited in NCBI’s Gene Expression and are accessible through GEO Series accession no. GSE155354 and no. GSE164161, respectively

## Supplementary materials and methods

### Immunohistochemical staining

Paraffin blocks were obtained from tissues and PDOs, previous fixation in 4% formaldehyde overnight at room temperature. Paraffin-embedded sections of 4 μm, for tissues, and 2.5 μm, for PDOs, were de-paraffinized, rehydrated and endogenous peroxidase activity was quenched (20 min, 1.5% H_2_O_2_). EDTA or citrate< based antigen retrieval was used depending on the primary antibody used. All primary antibodies were diluted in PBS containing 0.05% BSA, incubated overnight at 4 °C and developed with the Envision+ System HRP Labelled Polymer anti-Rabbit or anti-Mouse and 3,3′- diaminobenzidine (DAB). Samples were mounted in DPX and images were obtained with an Olympus BX61 microscope.

### Immunofluorescence analysis

For tissues and PDOs, the same protocol as IHC was followed. However, the samples were developed with Tyramide Signal Amplification System (TSA) and mounted in DAPI Fluoromount-G. Images were taken in an SP5 upright confocal microscope (Leica).

### Hematoxylin and eosin staining

Previously de-paraffinized sections were incubated with hematoxylin 30 s, tap water 5 min, 80% ethanol 0.15% HCl 30 s, water 30 s, 30% ammonia water (NH3(aq)) 30 s, water 30 s, 96% ethanol 5 min, eosin 3 s, and absolute ethanol 1 min. Samples were dehydrated, mounted in DPX, and images were obtained with an Olympus BX61 microscope.

### FISH

Fluorescent in-situ hybridization (FISH) analyses from control and IC_30_-treated PDOs were performed using commercial probes (Abbott Molecular Inc, Des Plaines, IL, USA), one including the centromeric alfa-satellite region specific for chromosome 8, and a second one containing locus-specific probes from the long arm of chromosome 13 and 21.

In brief, we performed a cytospin to concentrate nuclei in the FISH slide. Slides were pre-treated with pepsin for 5 minutes at 37°C. Samples and probe were co-denaturated at 80°C for five minutes and hybridized overnight at 37°C in a hot plate (Hybrite chamber, Abbot Molecular Inc.). Post-hybridization washes were performed at 73°C in 2x<u>s</u>odium <u>s</u>alt <u>c</u>itrate buffer (SSC) and at room temperature in 2xSSC, 0.1% NP-40 solution. Samples were counterstained with 4,6-diamino-2-phenilindole (DAPI)(Abbott Molecular Inc, Des Plaines, IL, USA). Results were analyzed in a fluorescence microscope (Olympus, BX51) using the Cytovision software (Applied Imaging, Santa Clara, CA). A minimum of 50 nuclei per case was analyzed.

### Comet assay

Comet assays were performed using Comet Assay Kit following manufacturer’s instructions. Pictures were taken using a Nikon Eclipse Ni-E epifluorescence microscope and tail moment was calculated using the OPENCOMET plugin for Fiji.

### Annexin V binding assay

Annexin V binding was determined by flow cytometry using the standard Annexin V Apoptosis Detection Kit APC. Single cells of treated PDOs with indicated combinations of 5-FU+Iri. were obtained and stained according to the manufacturer’s instructions, with Propidium Iodide staining for the DNA content. The cells were analyzed in the Fortessa analyzer.

### PDO initiating capacity assay

For PDO Initiating Capacity assay, 300 or 600 single PDO cells were plated in 96-well plates in 10 μL Matrigel. After 11 days in culture, the number of PDOs in each well was counted, photographs were taken for PDO diameter determination and cell viability was measured.

### Chromatin-immunoprecipitation assay (ChIP)

Control and IC20-treated PDOs were subjected to ChIP following standard procedures. Briefly, PDO cells were extracted with formaldehyde crosslinked for 10 min at room temperature and lysed for 20 min on ice with 500 μL of H_2_O plus 10 mM Tris-HCl pH 8.0, 0.25% Triton X-100, 10 mM EDTA, 0.5 mM EGTA, 20 mM β-glycerol-phosphate, 100 mM NA-orthovanadate, 10 mM NaButyrate and complete protease inhibitor cocktail. The supernatants were sonicated, centrifuged at 13.000 rpm for 15 min, and supernatants were incubated overnight with anti-p53 antibody in RIPA buffer. Precipitates were captured with 35 mL of protein A-Sepharose, extensively washed and analysed by ChIP-qPCR. Primers used are listed in Supplementary Table S7. Inputs were used to normalize the ChIP-qPCR and samples were compared to control IgGs.

### PDOs infection

hFLiG plasmid was used for in vivo detection of metastasis, H2BeGFP plasmid was used for flow cytometry experiments and lentiCRISPR v2 was used for knock-out experiments. Three sgRNA against *TP53* gene were designed using Benchling (Supplementary Table S7). Lentiviral production was performed transfecting in HEK293T cells the lentiviral vectors and the plasmid of interest. One day after transfection, medium was changed, and viral particles were collected 24 hours later and then concentrated using Lenti-X Concentrator. PDOs were infected by resuspending single cells in concentrated virus diluted in complete medium, centrifuged for 1 h at 650 rcf, and incubated for 5 hours at 37°C. Cells were then washed in complete culture medium and seeded as described above.

### Description of the patient gene expression data sets

Transcriptomic and available clinical data datasets from colorectal, breast, prostate and lung cancer were downloaded from the open-access resource CANCERTOOL. For CRC we used the Marisa (GSE39582) data set, which includes expression and clinical data for 566 patients with CRC and 19 non-tumoral colorectal mucosa, and the Jorissen (GSE14333) data set and the TCGA data set with expression and clinical data of 226 and 329 CRC patients, respectively. For Lung cancer we used the Okayama (GSE31210) data set, which includes expression profiles of 226 lung adenocarcinomas and the TCGA data set with 434 patients.

### Association of the signatures with clinical outcome

Association of the signatures expression with relapse was assessed in the cancer transcriptomic data sets using a Kaplan-Meier estimates and Cox proportional hazard models. A standard log-rank test was applied to assess significance between groups. This test was selected because it assumes the randomness of the possible censorship. All the survival analyses and graphs were performed with R using the survival (v.3.2-3) and survimer (v.0.4.8) packages and a p-value<0.05 was considered statistically significant (see Supplementary Table S4).

### Supplementary figures and tables

Fig. S1. Low-dose CT treatment induces a quiescent-like state to CRC PDO in the absence of persistent DNA damage and senescence.

Fig. S2. Low-dose CT induces a robust p53 signaling.

Fig. S3. p53 and p21 dependency of the feISC signature.

Fig. S4. TQL cells retain tumor initiating capacity.

Fig. S5. Identification of a fetal ISC signature with prognostic value in cancer.

Fig. S6. Acquisition of quiescent phenotype by CT treatment in patients.

Table S1. Patient-derived organoids used in this study.

Table S2. Differentially expressed genes between IC20 or IC30 and control PDOs.

Table S3. Expression correlation matrix from CT induced feISC genes in the Marisa (Marisa et al., 2013) dataset.

Table S4. Cox proportional hazards analysis of the feISC signature.

Table S5. Positive correlation of individual genes to the rest of the cohort.

Table S6. Human gastrointestinal tumor samples used in this study.

Table S7. List of oligonucleotides for RT-qPCR and ChIP-qPCR and sgRNA for CIRSPR/Cas9 knockout used in this study.

Table S8. Materials table.

**Figure S1.**
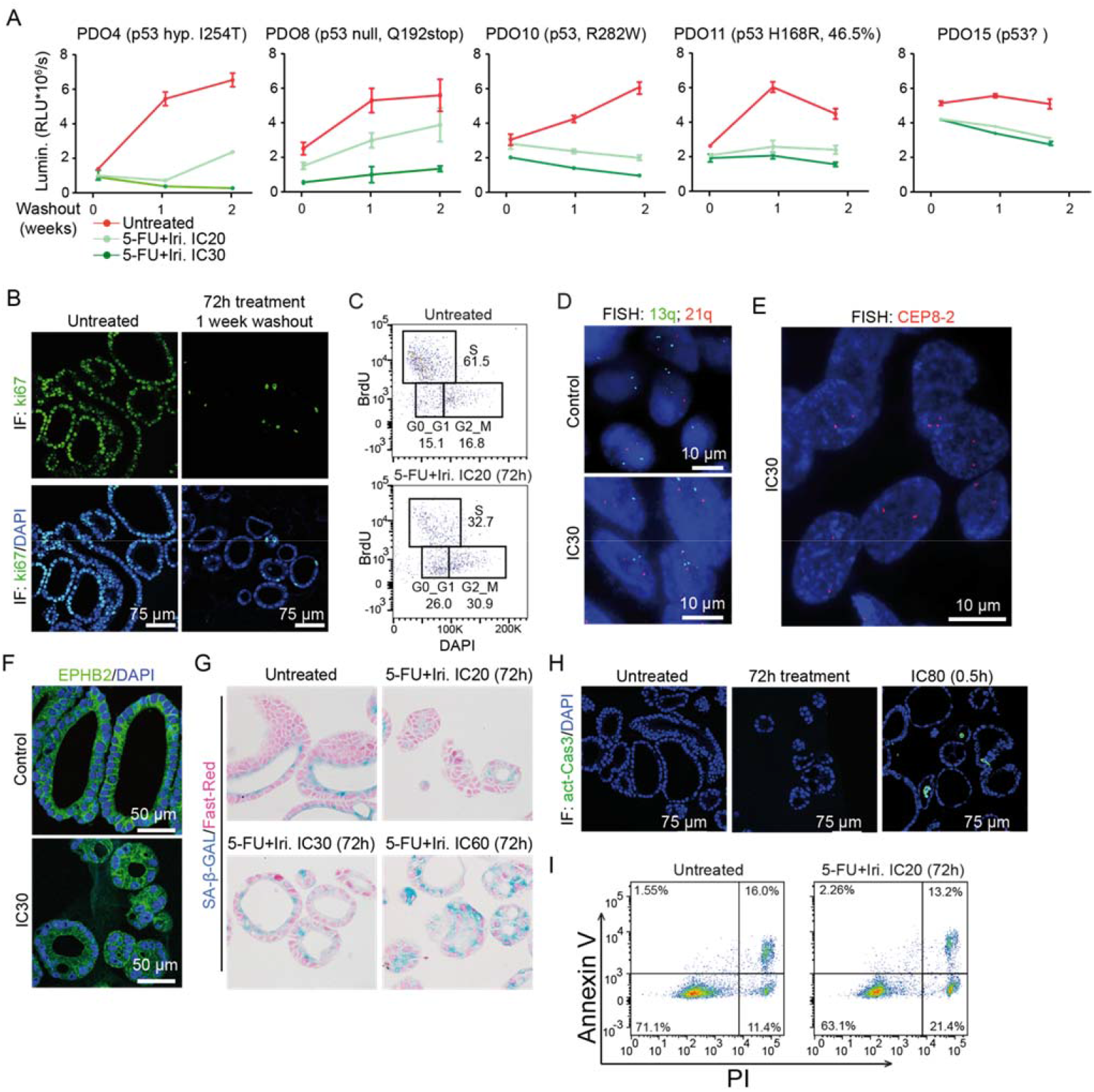
Related to Figure 1. Low-dose CT treatment induces a quiescent-like state to CRC PDO in the absence of persistent DNA damage and senescence. **(A)** Quantification of cell viability of the different PDOs untreated or pretreated with 5- FU+Iri. for 72 hours and then maintained in fresh medium for 1 and 2 weeks after washout. **(B)** Representative images of the proliferation marker ki67 staining by IF in PDO5 tumoroid treated with 5-FU+Iri. at IC_20_ for 72 hours and after being maintained in fresh medium for 1 week. **(C)** Flow cytometry analysis showing BrdU incorporation of PDO5 after 72 hours of 5-FU+Iri. treatment, compared with the control. Three boxes are shown, representing cells in G_0_/G_1_, S and G_2_/M cell cycle, respectively. **(D and E)** Representative images of fluorescent in-situ hybridization (FISH) analysis from control and IC_30_-treated PDO5 using probes for **(D)** 13q (green) and 21q (red) and **(E)** the centromeric probe CEP8-2 (red). **(F)** Representative images of IF analysis using the surface marker EPHB2 in control and IC_30_-treated PDO5 tumoroid. DAPI is used as a nuclear marker. **(G)** Analysis of SA-β-Gal activity in PDO5 cells treated with 5-FU+Iri. as indicated for 72 hours. Representative images were obtained with Olympus BX61. **(H)** Representative IF images of cleaved-caspase 3 staining in PDO5 treated with 5- FU+Iri. at IC_20_ at the indicated time points and with IC_80_ as a positive control. **(I)** Cytometry analysis of Annexin V binding in PDO5 untreated or treated as indicated. SA-β-Gal, SA-β-Galactosidase; 5-FU, 5-fluorouracil; Iri, irinotecan; CT, control; IC_20_ and IC_30_, 5-FU+Iri. treatment that results in 20% and 30% cell death, respectively, compared with untreated cell growth.

**Figure S2.**
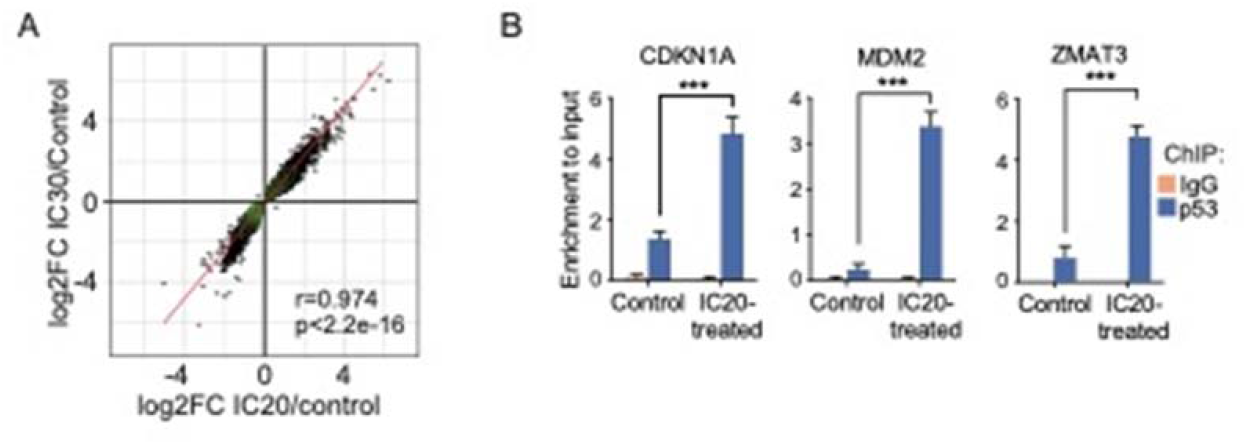
Related to Figure 2. Low-dose CT induces a robust p53 signaling. **(A)** Linear association of the genes differentially expressed in treated PDO5 compared with the control. Dots represent the log_2_ fold change values of genes for IC_20_ compared with control (x-axis) and IC_30_ compared with control (y-axis). The Pearson correlation and p value are shown. **(B)** ChIP-qPCR analysis of p53 binding in untreated and IC_20_- treated PDO5 in a subset of putative p53 target genes expressed as relative enrichment normalized to the input. The statistical analysis in (B) was performed by T-Student test, comparing treated with untreated condition. p values are indicated as ****p < 0.0001. CT, IC_20_ and IC_30_, 5- FU+Iri. treatment that results in 20% and 30% cell death, respectively, compared with untreated cell growth.

**Figure S3.**
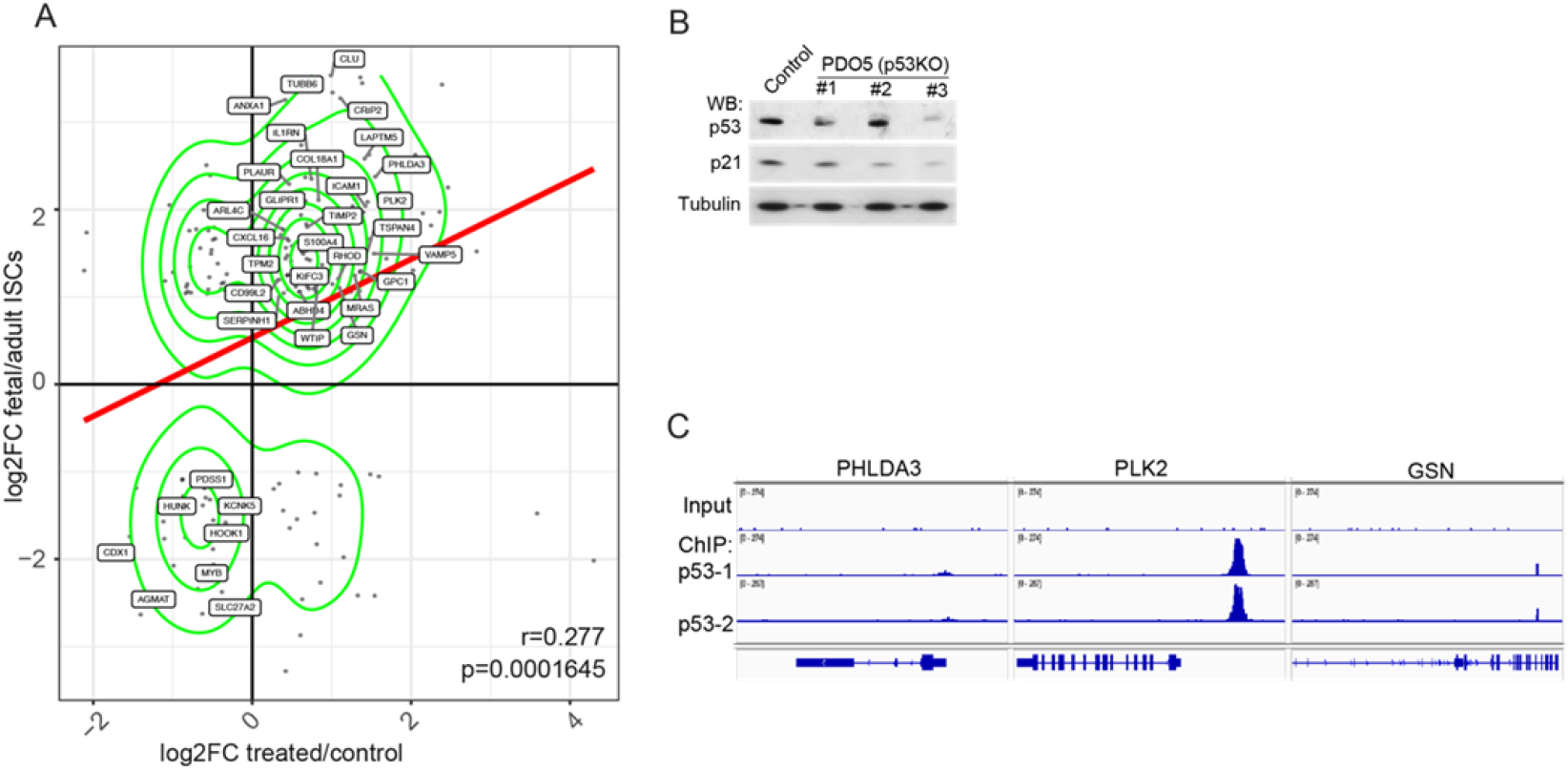
Related to Figure 3. p53 and p21 dependency of the feISC signature. **(A)** Scatter plot and linear regression line of the genes differentially expressed between CT treated and control PDOs and fetal compared with adult intestinal stem cell. Dots represent the log2 fold change values of genes for treated versus control (xaxis) and fetal versus adult intestinal stem cell (y-axis). The Pearson correlation and p value are shown. Genes included in the 28up+8down-feISC signature are indicated. **(B)** WB analysis of p53 levels and its downstream target p21 in CRISPR-Cas9-engineered p53 KO pools. **(C)** Representation of some 28up-feISC genes distribution in the indicated genomic regions obtained from ChIP-sequencing analysis in IC_20_-treated PDO5 (n=2).

**Figure S4.**
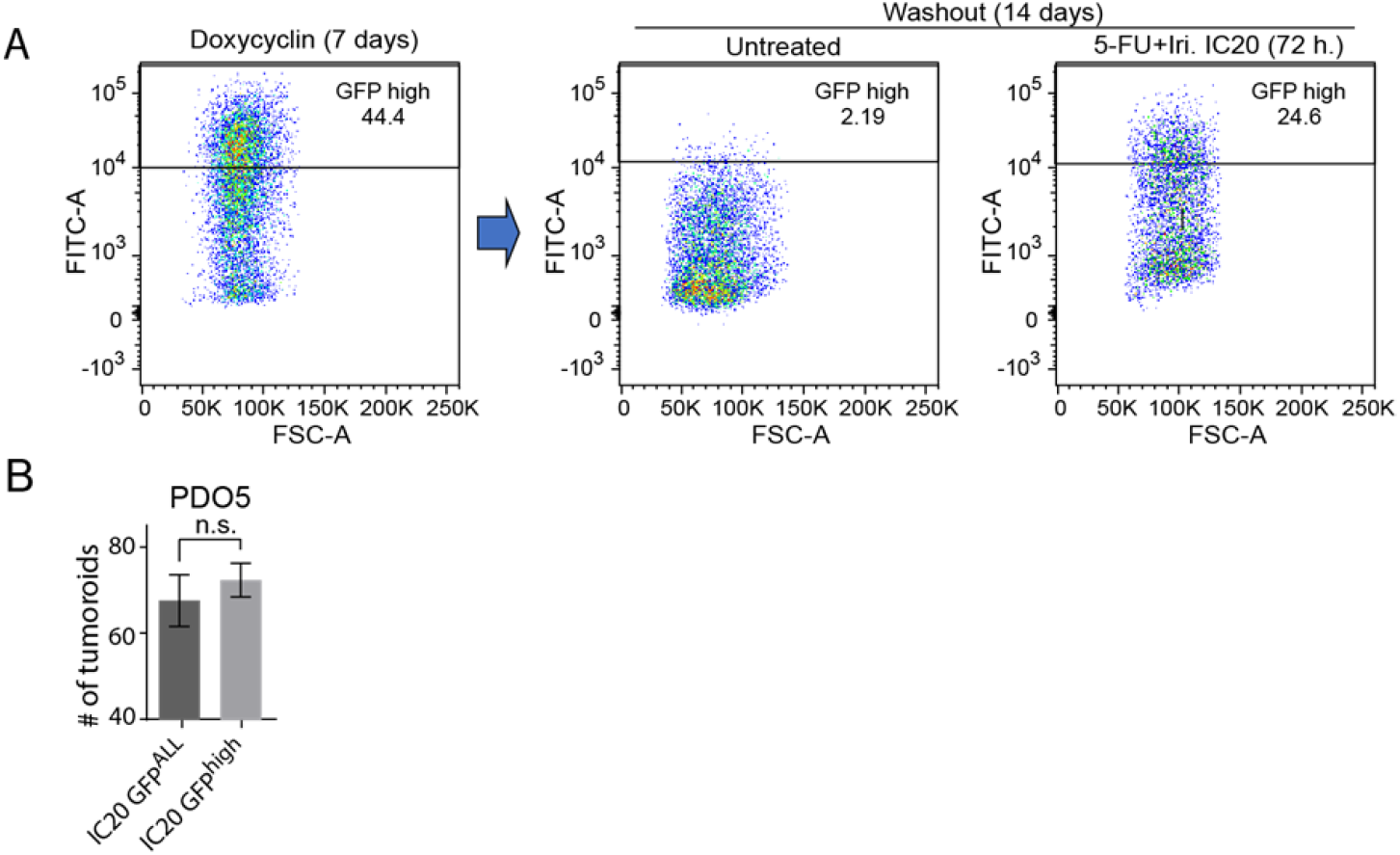
Related to Figure 4. TQL cells retain tumor initiating capacity. **(A)** Analysis of GFP distribution by flow cytometry of PDO5 cells carrying a doxycycline-inducible GFP-H2B construct. Cells were treated for 6 days with doxycycline to induce GFP-H2B expression and then left untreated or treated with 5- FU+Iri. IC_30_ for 72 hours and maintained in fresh medium for 2 additional weeks. Quiescent cells that retained high or low GFP levels were purified by cell sorting. **(B**) Number of PDOs generated from seeding 300 GFP^high+low^ and GFP^high^ sorted cells after 2 weeks with fresh medium. n.s., no significant; IC_20_, 5-FU+Iri. treatment that results in 20% cell death, compared with untreated cell growth.

**Figure S5.**
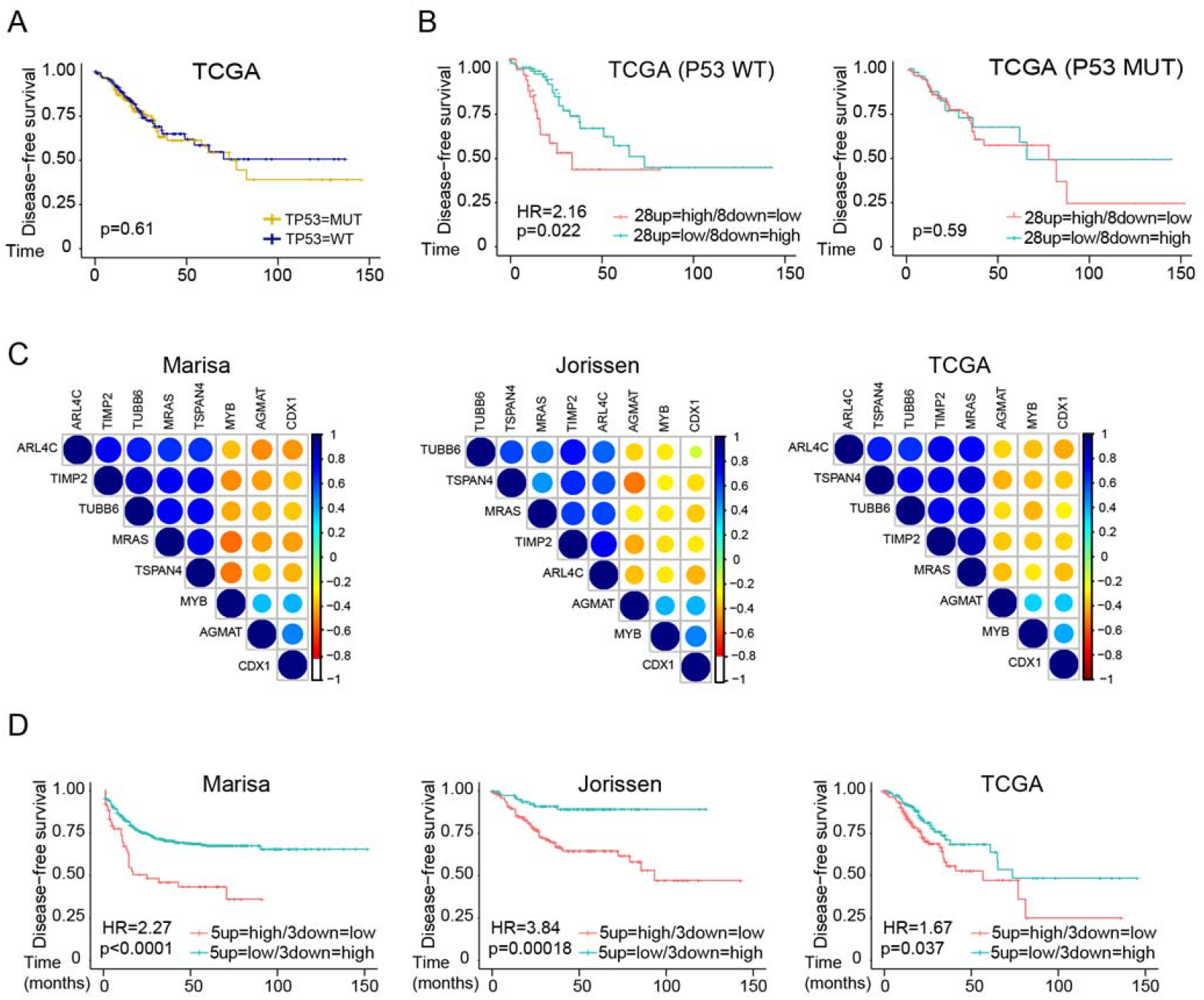
Related to Figure 5. Identification of a fetal ISC signature with prognostic value in cancer. (**A**) Kaplan-Meier representation of disease-free survival probability over time of patients classified according to their *TP53* status (*TP53* WT n=144 and *TP53* MUT n=177) in the TCGA colorectal cancer dataset. **(B)** Kaplan-Meier representation of disease-free survival probability over time of patients, from the TCGA dataset, classified according to their cluster analysis of the 28up+8down-feISC signature (data not shown) for patient groups from *TP53* WT (28up=high/8down=low *n=*47 and 28up=low/8down=high *n=*97) and *TP53* mutant (28up=high/8down=low *n=*128 and 28up=low/8down=high *n=*49). **(C)** Expression correlation matrix according to the 5up+3down-feISC signature in the Marisa, Jorissen and TCGA datasets. Positive and negative correlation is shown in blue and red, respectively. The size of circles and color intensity are proportional to the Pearson correlation coefficient found for each gene pair. **(D)** Kaplan–Meier curves of disease-free survival probability over time of patients classified according to their cluster analysis of the 28up+8down-feISC signature (data not shown), for Marisa (5up=high/3down=low *n=*59 and 5up=low/3down=high *n=*507), Jorissen (5up=high/3down=low *n=*137 and 5up=low/3down=high *n=*89) and TCGA (5up=high/3down=low *n=*128 and 5up=low/3down=high *n=*105) colorectal databases. For statistical analysis of the Kaplan-Meier estimates we used Cox proportional hazards models (See Supplementary Table S4). HR, hazard ratio.; p, p-value.

**Figure S6.**
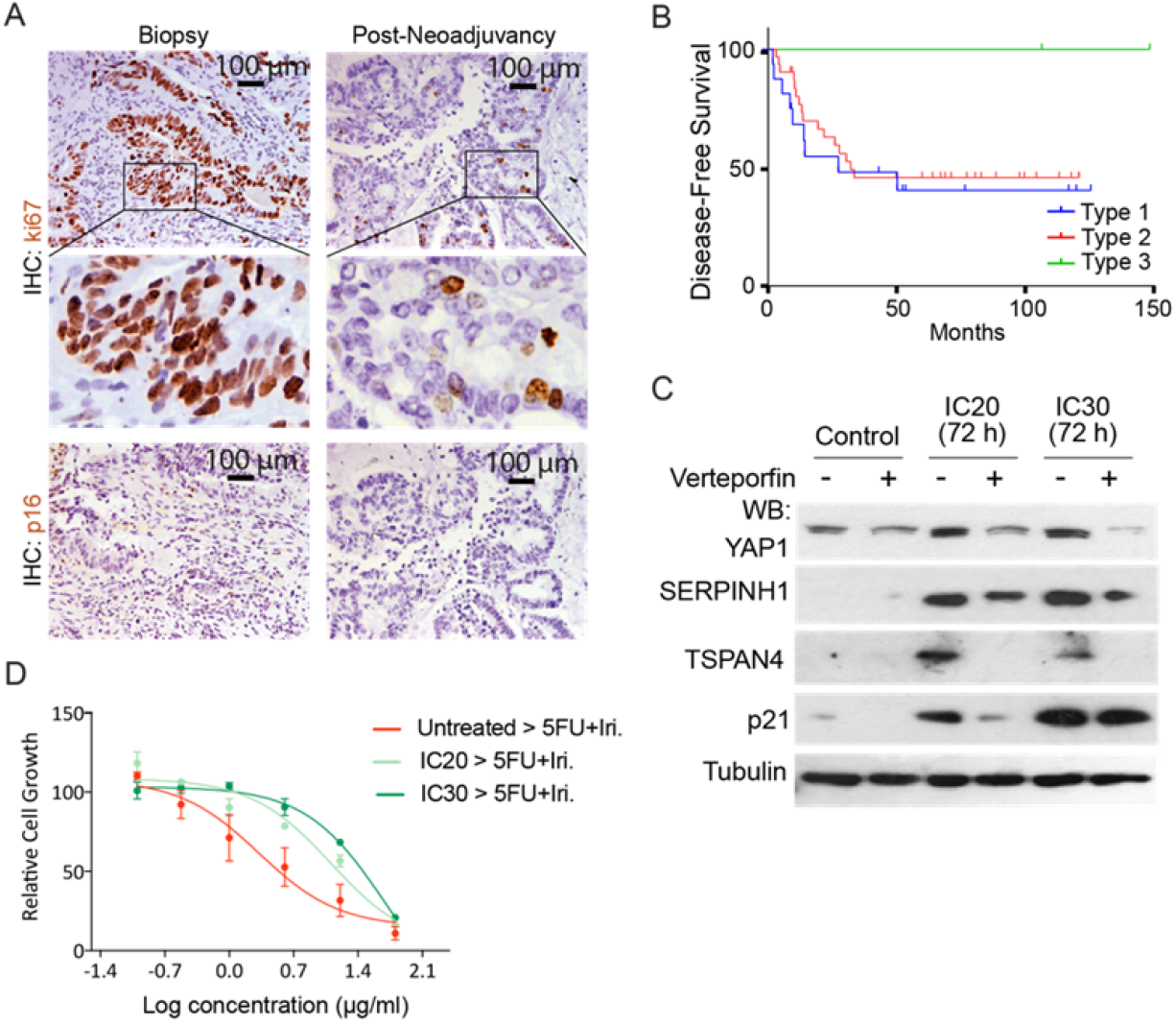
Related to Figure 6. Acquisition of quiescent phenotype by CT treatment in patients. **(A)** IHC analysis of Ki67 and p16 in representative type 2 colorectal tumor samples (Table S6 #17) from the same patient at diagnosis (biopsy) and after neoadjuvant therapy at the time of surgery (post-neoadjuvancy). (**B**) Disease-free survival analysis (Kaplan-Meier curves) of patients stratified according to ki67- related tumor type. **(C)** WB analysis of control and treated *TP53* WT Ls174T CRC cells collected after 24 hours of 5-FU+Iri. treatment alone or in combination with the YAP1 inhibitor verteporfin at a final concentration of 5 < M. **(D)** Dose-response assay of PDO5 cells untreated or previously treated for 72 hours with IC_20_-IC_30_ 5-FU+Iri.

**Supplementary Table S1.** Patient-derived organoids used in this study. The mutations and the corresponding chemotherapy concentrations that reduce a 20 and 30% of the cell growth (IC20 and IC30, respectively) are indicated for each PDO.

| PDO | Mutations | IC20 (µg/mL) | IC30 (µg/mL) |
| --- | --- | --- | --- |
| PDO4 | TP53 I254T (100%)<br>EGFR S464L (97.21%) | 5-FU 1.25<br>Iri. 0.50 | 5-FU 2.00<br>Iri. 0.80 |
| PDO5 | KRAS G12D (66.43%) | 5-FU 0.14<br>Iri.] 0.06 | 5-FU 0.25<br>Iri. 0.10 |
| PDO8 | TP53 Q192stop (98.46%)<br>KRAS G13C (67.27%) | 5-FU 0.78<br>Iri. 0.31 | 5-FU 1.56<br>Iri. 0.63 |
| PDO10 | TP53 R282W (99.82)<br>FGFR2 C809W (62.72%)<br>KRAS A146V (80.25%) | 5-FU 6.25<br>Iri. 2.50 | 5-FU 12.50<br>Iri. 5.00 |
| PDO11 | TP53 H168R (46.54%)<br>FGFR2 N194stop (45.87%)<br>KRAS G12D (49.58%)<br>ERBB2 A87T (4.76%)<br>PIK3CA S874N (41.93%)<br>PDGFRA R293H (5.97%)<br>EGFR K960R (46.3%)<br>BRAF E71D (42.08%) | 5-FU 0.78<br>Iri. 0.31 | 5-FU 1.56<br>Iri. 0.63 |
| PDO15 | TP53 G262V (98.82%) | 5-FU 1.56<br>Iri. 0.63 | 5-FU 3.13<br>Iri. 1.25 |
| PDO66 | NRAS (G12S) (99.05%)<br>APC (S1110stop) (99.80) | 5-FU 0.63<br>Iri. 0.30 | 5-FU 2.50<br>Iri. 1.00 |

**Table S4.**
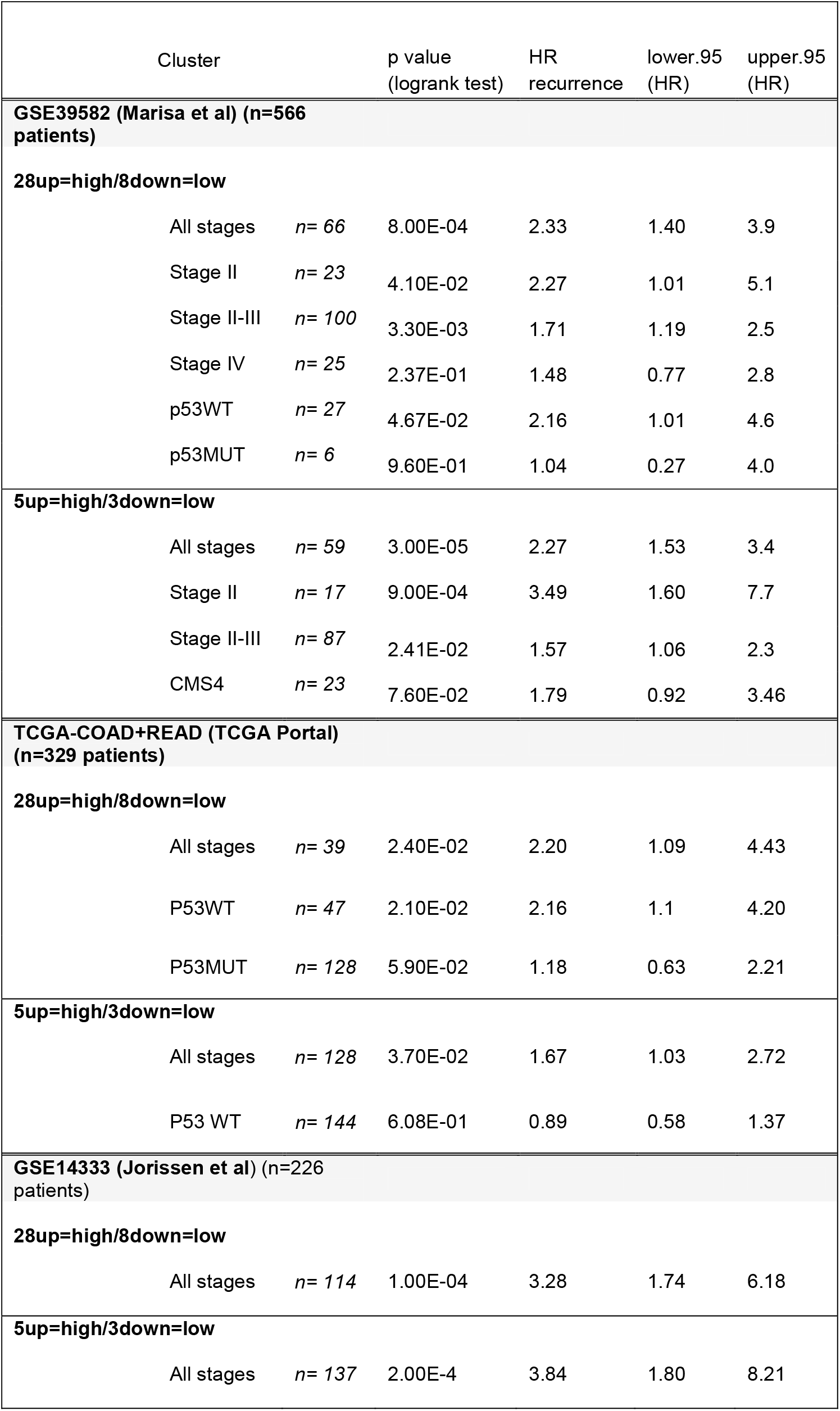
Cox proportional hazards analysis of the feISC signature. Association of the signature with recurrence-disease free survival. Related to Figure 4 and Figure S4.

**Table S5.** Positive correlation of individual genes to the rest of the cohort. Genes within the 5up+3down-feISC signature are highlighted in gray. Related to Figure 5 and S5.

| Gene | State | GSE39582<br>(Marisa et al) | TCGA-COAD+READ<br>(TCGA Portal) | GSE14333<br>(Jorissen et al) | Final Score |
| --- | --- | --- | --- | --- | --- |
| ABHD4 | up | 9,69 | 9,39 | 7,31 | 26,39 |
| ANXA1 | up | 9,44 | 11,85 | 8,82 | 30,12 |
| ARL4C | up | 13,77 | 14,96 | 12,84 | 41,57 |
| CD99L2 | up | 9,48 | 9,65 | 4,87 | 23,99 |
| CLU | up | 10,84 | 9,07 | 9,18 | 29,09 |
| COL18A1 | up | 14,22 | 14,74 | 10,80 | 39,76 |
| CRIP2 | up | 13,76 | 14,17 | 9,11 | 37,04 |
| CXCL16 | up | 8,12 | 7,54 | 8,59 | 24,25 |
| GLIPR1 | up | 8,08 | 9,84 | 9,18 | 27,10 |
| GPC1 | up | 11,18 | 11,40 | 8,45 | 31,03 |
| GSN | up | 9,85 | 11,30 | 3,41 | 24,56 |
| ICAM1 | up | 12,82 | 13,66 | 9,16 | 35,64 |
| IL1RN | up | 9,70 | 9,66 | 7,92 | 27,28 |
| KIFC3 | up | 10,77 | 14,54 | 3,25 | 28,57 |
| LAPTM5 | up | 13,56 | 14,90 | 11,43 | 39,88 |
| MRAS | up | 15,43 | 16,61 | 10,53 | 42,57 |
| PHLDA3 | up | 11,92 | 12,29 | 5,69 | 29,90 |
| PLAUR | up | 11,48 | 10,39 | 10,19 | 32,07 |
| PLK2 | up | 9,78 | 10,16 | 8,90 | 28,84 |
| RHOD | up | 8,63 | 4,08 | 7,79 | 20,50 |
| S100A4 | up | 11,56 | 10,19 | 9,35 | 31,09 |
| SERPINH1 | up | 13,66 | 13,66 | 10,22 | 37,53 |
| TIMP2 | up | 15,72 | 16,68 | 13,80 | 46,21 |
| TPM2 | up | 11,24 | 13,25 | 8,91 | 33,41 |
| TSPAN4 | up | 15,69 | 16,26 | 12,19 | 44,15 |
| TUBB6 | up | 15,37 | 15,78 | 11,66 | 42,81 |
| VAMP5 | up | 13,26 | 14,07 | 7,73 | 35,06 |
| WTIP | up | 8,59 | 12,75 | 5,23 | 26,57 |
| AGMAT | down | 3,74 | 3,21 | 3,32 | 10,28 |
| CDX1 | down | 3,61 | 3,13 | 3,37 | 10,12 |
| HOOK1 | down | 3,42 | 2,93 | 3,12 | 9,47 |
| HUNK | down | 2,73 | 2,72 | 2,74 | 8,18 |
| KCNK5 | down | 2,60 | 2,39 | 2,49 | 7,48 |
| MYB | down | 3,62 | 3,46 | 3,50 | 10,58 |
| PDSS1 | down | 3,46 | 2,98 | 2,42 | 8,86 |
| SLC27A2 | down | 2,69 | 2,12 | 2,22 | 7,03 |

**Supplementary Table S6.** Human gastrointestinal tumor samples used in this study. Paired samples at diagnosis (biopsy) and after neoadjuvant therapy at the time of surgery (postQ). The corresponding clinical data, ki67 subtype classification and presence of nuclear YAP1 is indicated.

| Patient nº | Tumor Localization | Clinical TNM | Treatment | ki67% postQ | ki67% biopsy | ki67 subtype | nuclear YAP1% postQ | nuclear YAP1% biopsy | OS (mo) | DFS (mo) |
| --- | --- | --- | --- | --- | --- | --- | --- | --- | --- | --- |
| 1 | Gastric | T4N1M1 | Chemotherapy | 20 | 40 | 2 | 30 | 0 | 7,13 | 5,40 |
| 2 | Colorectal | T3N1M1 | Chemotherapy + Targeted therapy | 1 | 55 | 3 | N/A | N/A | 115,23 | 115,23 |
| 3 | Gastric | T3N1 | Chemotherapy | 10 | 70 | 2 | 90 | 5 | 60,90 | 60,90 |
| 4 | Colorectal | T3N1M1 | Radiotherapy | 5 | 70 | 2 | N/A | N/A | 17,33 | 9,50 |
| 5 | Gastric | T3N1M0 | Chemotherapy | 5 | 60 | 2 | 80 | 5 | 26,73 | 12,23 |
| 6 | Colorectal | T3N0 | Chemotherapy | 30 | 70 | 2 | 50 | 2 | 53,10 | 53,10 |
| 7 | Gastric | T3N1M1 | Chemotherapy | 70 | 90 | 2 | 90 | 80 | 41,10 | 23,07 |
| 8 | Gastric | T3N1M1 | Chemotherapy + Targeted therapy | 10 | 5 | 1 | 70 | 0 | 89,83 | 40,23 |
| 9 | Colorectal | T3N1 | Chemotherapy | 15 | 75 | 2 | N/A | N/A | 94,23 | 94,23 |
| 10 | Gastric | T3N0M0 | Chemotherapy | 85 | 95 | 1 | 5 | 10 | 91,10 | 91,10 |
| 11 | Gastric | T2-3N1M0 | Chemotherapy | 90 | 90 | 1 | N/A | N/A | 93,37 | 93,37 |
| 12 | Gastric | T3N1M0 | Chemotherapy | 80 | 90 | 1 | N/A | N/A | 97,70 | 97,70 |
| 13 | Gastric | T3N0M0 | Chemotherapy | 20 | 70 | 2 | 0 | 0 | 25,23 | 9,67 |
| 14 | Colorectal | T4N1 | Chemotherapy | 80 | 60 | 1 | 95 | 20 | 38,17 | 12,43 |
| 15 | Gastric | T3N1M1 | Chemotherapy + Targeted therapy | 70 | 85 | 1 | 50 | 15 | 93,37 | 93,37 |
| 16 | Colorectal | T3N1 | Chemotherapy + Targeted therapy | 10 | 80 | 2 | 90 | 0 | 42,67 | 21,67 |
| 17 | Colorectal | T4N1M0 | Chemotherapy | 20 | 95 | 2 | 60 | 20 | 39,90 | 39,90 |
| 18 | Colorectal | T3N0M0 | Chemotherapy | 40 | 90 | 2 | N/A | N/A | 81,53 | 81,53 |
| 19 | Colorectal | T2N1M0 | Chemotherapy | 5 | 25 | 3 | 90 | 3 | 83,13 | 83,13 |
| 20 | Colorectal | T4N1M0 | Chemotherapy | 20 | 60 | 2 | 70 | 60 | 94,03 | 94,03 |
| 21 | Colorectal | T3N1M0 | Chemotherapy | 25 | 80 | 2 | N/A | 30 | 91,90 | 91,90 |
| 22 | Colorectal | T3N1M0 | Chemotherapy | 1 | 95 | 2 | N/A | N/A | 88,30 | 88,30 |
| 23 | Colorectal | T3N2bM0 | Chemotherapy | 1 | 90 | 2 | 30 | 30 | 76,53 | 76,53 |
| 24 | Gastric | T3N1M0 | Chemotherapy | 20 | 20 | 1 | 90 | 0 | 29,87 | 12,77 |
| 25 | Colorectal | T3N1M0 | Chemotherapy | N/A | N/A | N/A | N/A | N/A | 79,77 | N/A |
| 26 | Colorectal | T3N0M0 | Chemotherapy | N/A | N/A | N/A | N/A | N/A | 53,63 | N/A |
| 27 | Colorectal | T3N0M0 | Chemotherapy | 1 | 55 | 2 | 5 | 10 | 8,53 | 8,53 |
| 28 | Colorectal | T3N1M0 | Chemotherapy + Targeted | 40 | 65 | 2 | 30 | 1 | 121,23 | 27,47 |

| therapy |  |  |  |  |  |  |  |  |  |  |
| --- | --- | --- | --- | --- | --- | --- | --- | --- | --- | --- |
| 29 | Colorectal | T3N1M0 | Chemotherapy | N/A | N/A | N/A | N/A | N/A | 33,90 | 20,83 |
| 30 | Colorectal | T3N0 | Chemotherapy | 20 | 10 | 1 | 70 | 3 | 60,30 | 60,30 |
| 31 | Gastric | T3N0 | Chemotherapy | 20 | 30 | 1 | 100 | 0 | 12,17 | 9,10 |
| 32 | Gastric | T3N1 | Chemotherapy | 30 | 40 | 1 | N/A | N/A | 16,53 | 8,40 |
| 33 | Gastric | T3N0M0 | Chemotherapy | 30 | 70 | 2 | 80 | 5 | 77,93 | 77,93 |
| 34 | Colorectal | T3N1M1 | Chemotherapy<br>+ Targeted<br>therapy | 5 | 80 | 2 | 90 | 0 | N/A | N/A |
| 35 | Colorectal | T3N1M0 | Radiotherapy | 1 | 60 | 2 | 100 | 15 | 69,50 | 69,50 |
| 36 | Gastric | T3N1M0 | Chemotherapy | 25 | N/A | N/A | N/A | N/A | 39,37 | 39,37 |
| 37 | Gastric | T3N1M0 | Chemotherapy | 20 | 55 | 2 | 2 | 2 | 65,10 | 65,10 |
| 38 | Gastric | T3N1M0 | Chemotherapy | 10 | 20 | 1 | 60 | 20 | 39,23 | 22,73 |
| 39 | Pancreas | T4N0M0 | Chemotherapy | 0 | N/A | N/A | N/A | N/A | 45,63 | 15,63 |
| 40 | Colorectal | T3N1M1 | Chemotherapy | 25 | 30 | 1 | 90 | 10 | 18,23 | 6,00 |
| 41 | Colorectal | T3N0M0 | Radiotherapy | 25 | 40 | 1 | 30 | 0 | 41,77 | 41,77 |
| 42 | Colorectal | T3N0M1 | Chemotherapy<br>+ Targeted<br>therapy | 40 | 90 | 2 | 90 | 0 | 22,50 | 10,13 |
| 43 | Colorectal | T4N1M0 | Chemotherapy | 0 | 65 | 2 | 0 | 5 | 45,60 | 11,90 |
| 44 | Colorectal | T3N1 | Chemotherapy | 5 | 15 | 1 | 0 | 5 | 8,70 | 8,70 |
| 45 | Colorectal | T3N0M0 | Chemotherapy | 1 | 70 | 2 | N/A | N/A | 62,73 | 25,00 |
| 46 | Colorectal | T3N1M1 | Chemotherapy<br>+ Targeted<br>therapy | 5 | 50 | 2 | 80 | 0 | 33,47 | 18,43 |
| 47 | Colorectal | T3N2M0 | Chemotherapy | 60 | 95 | 2 | 0 | 2 | 59,63 | 26,40 |
| 48 | Colorectal | T3N1M0 | Chemotherapy | 20 | 60 | 2 | 30 | 0 | 56,20 | 56,20 |
| 49 | Gastric | T4N0M0 | Chemotherapy | 50 | 60 | 1 | 50 | 50 | 42,70 | 42,70 |
| 50 | Colorectal | T3N0M0 | Chemotherapy | 10 | 70 | 2 | 5 | 0 | 47,60 | 47,60 |
| 51 | Gastric | T2N1M0 | Chemotherapy | 15 | 75 | 2 | 2 | 0 | 38,73 | 16,70 |
| 52 | Colorectal | T3N0M0 | Chemotherapy | 5 | 25 | 2 | 30 | 0 | 63,37 | 63,37 |
| 53 | Colorectal | T3N1M0 | Chemotherapy | 20 | 95 | 2 | 30 | 0 | 9,00 | 9,00 |
| 54 | Gastric | T3N3M0 | Chemotherapy | 15 | 60 | 2 | 30 | 0 | 11,93 | 11,03 |
| 55 | Colorectal | T4N1M0 | Radiotherapy | 20 | 30 | 1 | 15 | 0 | 34,73 | 34,73 |
| 56 | Colorectal | T2N1M0 | Chemotherapy | N/A | N/A | N/A | N/A | N/A | 53,27 | N/A |
| 57 | Colorectal | T3N1M0 | Chemotherapy | 45 | 85 | 2 | 50 | 5 | 54,47 | 54,47 |
| 58 | Colorectal | T3N0M0 | Chemotherapy | 70 | 90 | 2 | 90 | 20 | 50,67 | 50,67 |
| 59 | Colorectal | T3N0M0 | Chemotherapy | 45 | 80 | 2 | 70 | 25 | 12,23 | 5,00 |
| 60 | Colorectal | T4N2M0 | Chemotherapy | 75 | 95 | 2 | 40 | 0 | 22,47 | 4,27 |
| 61 | Colorectal | T3Nx<br>(sigma)+<br>T2N0<br>(rectum)<br>M1 | Radiotherapy | 65 | 65 | 1 | N/A | N/A | 36,47 | 3,27 |
| 62 | Colorectal | T3N0<br>(rectum)<br>+ T4Nx<br>(bladder)<br>M1 | Radiotherapy | 80 | 95 | 1 | 80 | 5 | 5,37 | 3,53 |

**Supplementary Table S7.**
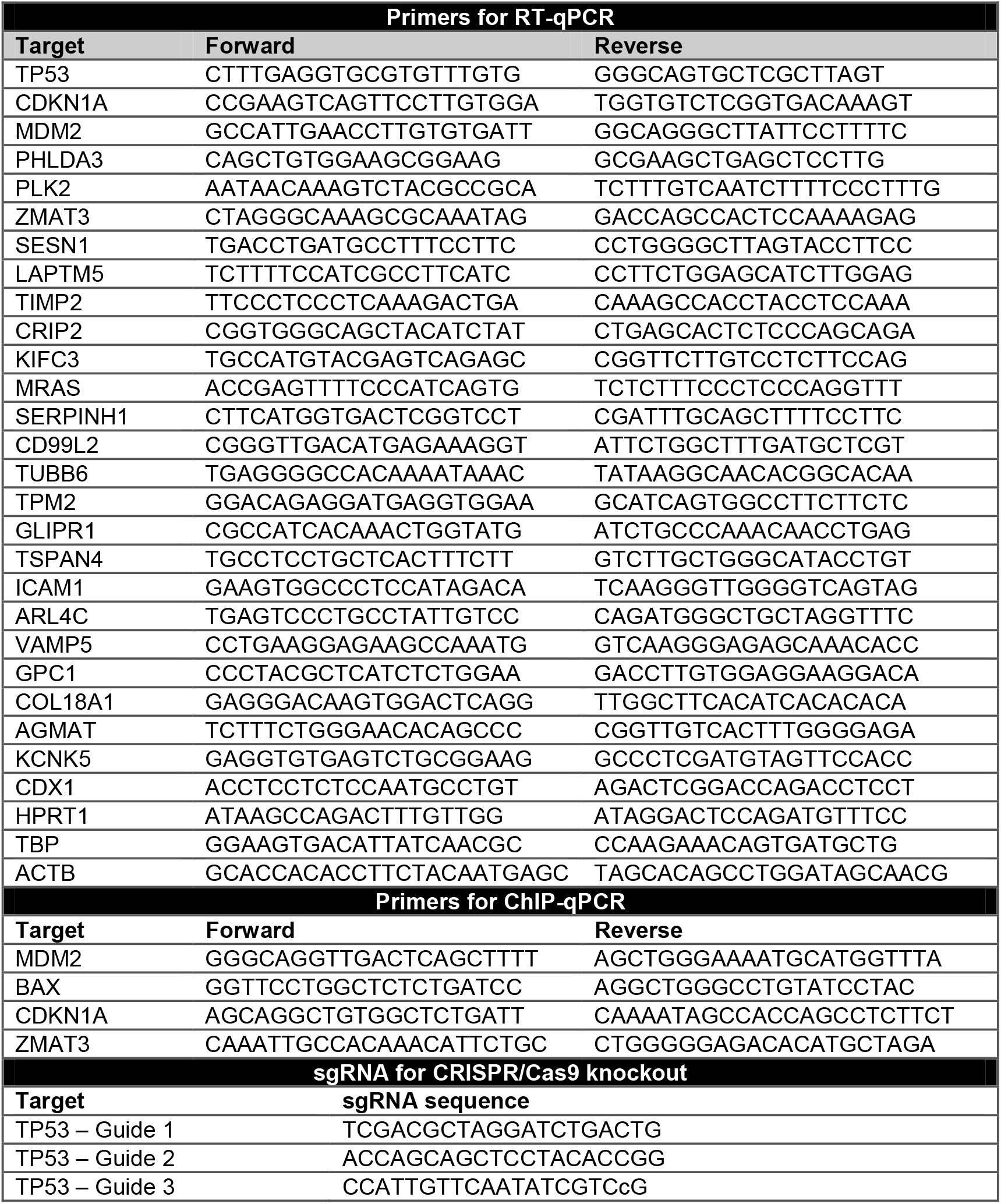
List of oligonucleotides for RT-qPCR and ChIP-qPCR and sgRNA for CIRSPR/Cas9 knockout used in this study.

**Table S8.** Materials table.

| REAGENT or | SOURCE | IDENTIFIER |
| --- | --- | --- |
| **Antibodies** |  |  |
| Mouse monoclonal anti-γH2AX (pS139) | BD Biosciences | Cat#564719; RRID:AB_2738913 |
| Mouse monoclonal anti-Ki67 (MM1) | Leica Biosystems | Cat#NCL-Ki67-MM1; RRID:AB_442101 |
| Rabbit polyclonal anti-Cleaved Caspase-3 (Asp175) | Cell Signaling | Cat#9661; RRID:AB_2341188 |
| Mouse monoclonal anti-p53 DO-1 | Abcam | Cat#ab1101; RRID:AB_297667 |
| Rabbit monoclonal anti-p21 [EPR362] | Abcam | Cat#ab109520; RRID:AB_10860537 |
| Rabbit monoclonal anti-CKN2A/p16INK4a [EPR1473] | Abcam | Cat#ab108349; RRID:AB_10858268 |
| Goat polyclonal anti-EphB2 | RD Systems | Cat#AF467; RRID:AB_355375 |
| Rabbit polyclonal anti-CD99L2 | Abcam | Cat#ab224164 |
| Mouse monoclonal anti-TIMP2 [3A4] | Abcam | Cat#ab1828; RRID:AB_2256129 |
| Rabbit polyclonal anti-MRas | Abcam | Cat#ab26303; RRID:AB_470849 |
| Anti-TUBB6 | Abcam | Cat#PA5-P8948 |
| Recombinant Anti-ICAM1 antibody [EPR4776] | Abcam | Cat#ab109361; RRID:AB_10958467 |
| Recombinant Anti-Hsp47 antibody [EPR4217] | Abcam | Cat#ab109117; RRID:AB_10888995 |
| Recombinant Anti-YAP1 antibody [EP1674Y] | Abcam | Cat#ab52771; RRID:AB_2219141 |
| TSPAN4 Polyclonal Antibody | Thermo Fisher Scientific | Cat#PA5-69344; RRID:AB_2688603 |
| Monoclonal Anti-S100A4 antibody produced in mouse | Atlas Antibodies | Cat#AMAB90599; RRID:AB_2665603 |
| Anti-Histone H3 antibody-Nuclear Marker and ChIP Grade | Abcam | Cat#ab791; RRID:AB_302613 |
| Anti-Histone H4 antibody–ChIP Grade | Abcam | Cat#ab10158; RRID:AB_296888 |
| Mouse monoclonal anti-alpha-Tubulin (B-5-1-2) | Sigma-Aldrich | Cat#T6074; RRID:AB_477582 |
| Goat Anti-Rabbit Immunoglobulins/HRP antibody (2ary) | Agilent | Cat#P0448; RRID:AB_2617138 |
| Rabbit Anti-Mouse Immunoglobulins/HRP antibody (2ary) | Agilent | Cat#P0260; RRID:AB_2636929 |
| Polyclonal Rabbit Anti-Goat Immunoglobulins/HRP antibody (2ary) | Agilent | Cat#P0449; RRID:AB_2617143 |
| **Biological Samples** |  |  |
| Patient-derived organoids (PDO): PDO4, PDO5, PDO8, PDO10, PDO11, PDO15 | Hospital del Mar (Barcelona) | MARBiobanc (https://marbiobanc.imim.es) |
| Patient-derived organoids (PDO): PDO66 | From Alberto Muñoz Lab (Fernández-Barral et al., 2020) | RetBioH (www.redbiobancos.es) |
| Human gastrointestinal tumors blocks | Hospital del Mar (Barcelona) | MARBiobanc (https://marbiobanc.imim.es) |
| **Chemicals, Peptides, and Recombinant Proteins** |  |  |
| Collagenase II from *Clostridium histolyticum* | Sigma-Aldrich | Cat#C6885 |
| Hyaluronidase from bovine testes | Sigma-Aldrich | Cat#H3506 |
| DMEM/F-12 Advanced | GIBCO | Cat#12634028 |
| Primocin | Invitrogen | Cat#ant-pm-1 |
| B-27 Supplement (50X) | GIBCO | Cat#17504044 |
| N-2 supplement (100X) | GIBCO | Cat#17502048 |
| Nicotinamide | Sigma-Aldrich | Cat#N3376 |
| N-Acetyl-L-cysteine | Sigma-Aldrich | Cat#A7250 |
| Recombinant Human Noggin | PeproTech | Cat#120-10C |
| Recombinant Human R-Spondin-1 | PeproTech | Cat#120-38 |
| Y-27632 dihydrochloride (ROCK inhibitor) | Sigma-Aldrich | Cat#Y0503 |
| Prostaglandin E2 | Tocris | Cat#2296 |
| SB 202190 | Sigma-Aldrich | Cat#S7067 |
| A8301 (ALK inhibitor) | Sigma-Aldrich | Cat#SML0788 |
| hEGF | Sigma-Aldrich | Cat#E9644 |
| Gastrin I (human) | Tocris | Cat#3006 |
| Corning Matrigel Basement Membrane Matrix, LDEV-free | Corning | Cat#354234 |
| 5-Fluorouracil (5-FU) | Accord Healthcare | Cat#606544.3 |
| Irinotecan | Accord Healthcare | Cat#713386.5 |
| Dasatinib | Selleckchem | Cat#S1021 |
| Verteporfin | Selleckchem | Cat#S1786 |
| D-Luciferin | Goldbio | Cat#LUCK |
| PhosSTOP phosphatase inhibitor cocktail | Roche | Cat#PHOSS-RO |
| cOmplete Mini protease inhibitor cocktail | Roche | Cat#11836170001 |
| DPX mountant | Sigma-Aldrich | Cat#06522 |
| DAPI Fluoromount-G | Southern Biotech | Cat#0100-20 |
| Protein A-Sepharose CL-4B | GE Healthcare | Cat#17-0780-01 |
| Protein G-Sepharose 4 Fast Flow | GE Healthcare | Cat#17-0618-01 |

| Critical Commercial Assays |  |  |
| --- | --- | --- |
| Dako Envision+ System-HRP Labelled Polymer anti-Rabbit | Agilent | Cat#K4003 |
| Envision+ System-HRP Labelled Polymer anti-Mouse | Agilent | Cat#K4001 |
| Dako Liquid DAB+ Substrate Chromogen System | Agilent | Cat#K3468 |
| TSA Plus Cyanine 3/Fluorescein System | PerkinElmer | Cat#NEL753001KT |
| EZ-ECL Chemiluminescence Detection Kit for HRP | Biological Industries | Cat#20-500-120 |
| ECL Prime Western Blotting System | GE Healthcare | Cat#RPN2232 |
| CellTiter-Glo Luminescent Cell Viability Assay | Promega | Cat#G7571 |
| APC BrdU Flow Kit | BD Biosciences | Cat#552598 |
| Senescence β-Galactosidase Staining Kit | Cell Signaling | Cat#9860S |
| Cell Event Senescence Green Flow Cytometry Assay Kit | Invitrogen | Cat#C10840 |
| CometAssay Kit | Trevigen | Cat#4250-050-K |
| RNeasy Micro Kit | Qiagen | Cat#74004 |
| RT-First Strand cDNA Synthesis Kit | GE Healthcare Life Sciences | Cat#27-9261-01 |
| SYBR Green I Master Kit | Roche | Cat#04887352001 |
| Annexin V Apoptosis Detection Kit APC | Invitrogen | Cat#88-8007 |
| Lenti-X Concentrator | Clontech | Cat#631232 |
| Experimental Models: Cell Lines |  |  |
| Human: HEK293T | ATCC | CRL-11268 |
| Human: HCT116 | ATCC | CCL-247 |
| Human: Ls174T | ATCC | CL-188 |
| Human: SW480 | ATCC | CCL-228 |
| Human: HT29 | ATCC | HTB-38D |
| **Experimental Models: Organisms/Strains** |  |  |
| Mouse: NU/J (*Foxn1<sup>mut</sup>*) | The Jackson Laboratory | JAX: 002019 |
| **Oligonucleotides** |  |  |
| Primers for RT-qPCR, see Table S7 | This study | N/A |
| Primers for Chip-qPCR, see Table S7 | This study | N/A |
| gRNA against *TP53*, see Table S7 | This study | N/A |
| **Recombinant DNA** |  |  |
| Plasmid: pMD2.G | From Trono Lab, unpublished | Addgene plasmid #12259 |
| Plasmid: pCMV-dR8.2 dvpr | Stewart et al., 2003 | Addgene plasmid #8455 |
| Plasmid: lentiCRISPR v2 | Sanjana et al., 2014 | Addgene plasmid #52961 |
| Plasmid: pLEX-hFLiG | Celià-Terrassa & Kang, 2018 | N/A |
| Plasmid: pLTPC-H2BeGFP | Gift from Héctor G. Palmer Lab, unpublished | N/A |
| **Software and Algorithms** |  |  |
| GraphPad Prism 6 | Graphpad Software | https://www.graphpad.com; RRID:SCR_002798 |
| Fiji (Image J) | Schneider et al., 2012 | https://www.fiji.sc; RRID: SCR_002285 |
| Benchling CRISPR design | Benchling | https://www.benchling.com; RRID: SCR_013955 |
| FlowJo 10.6.2 | BD Biosciences | https://www.flowjo.com; RRID: SCR_008520 |
| LightCycler Software | Roche | http://www.roche-applied-science.com/shop/products/absolute-quantification-with-the-lightcycler-carousel-based-system; RRID: SCR_012155 |
| Adobe Photoshop | Adobe Software | https://www.adobe.com/products/photoshop.html; RRID: SCR_014199 |
| RStudio | RStudio Team | https://rstudio.com/ |
| GSEA | Broad Institute | https://www.gsea-msigdb.org/gsea/index.jsp |
| ggplot | Bioconductor |  |
| Corrplot | CRAN | https://cran.r-project.org/web/packages/corrplot/index.html |
| Survminer | CRAN | https://CRAN.R-project.org/package=survminer |
| Survival | CRAN | https://CRAN.R-project.org/package=survival |
| Heatmaply | CRAN | https://CRAN.R-project.org/package=heatmaply |
| Pheatmap | CRAN | https://CRAN.R-project.org/package=pheatmap |
| Limma | Bioconductor | https://bioconductor.org/packages/release/bioc/html/limma.html |
| DESeq2 R package | Bioconductor | https://bioconductor.org/packages/release/bioc/html/DESeq2.html |
| TopHat | Kim et al 2013 | https://ccb.jhu.edu/software/tophat/index.shtml |
| HTSeq | Anders et al. 2015 | https://htseq.readthedocs.io/en/master/ |

